# The aminoglycoside streptomycin triggers ferroptosis in tumor initiating cells

**DOI:** 10.1101/2022.10.31.514281

**Authors:** Hélène Guillorit, Sébastien Relier, Benjamin Zagiel, Audrey Di Giorgio, Chantal Cazevieille, Lucile Bansard, Céline Bouclier, Xavier Mialhe, Morgan Brisset, Szimonetta Hideg, Armelle Choquet, Chris Planque, Amandine Bastide, Julie Pannequin, Maria Duca, Françoise Macari, Alexandre David

## Abstract

Compelling evidence suggests that tumor initiating cells (TIC) are the roots of current shortcomings in advanced and metastatic cancer treatment. TIC represents a minor subpopulation of tumor cells endowed with self-renewal and multi-lineage differentiation capacity, which can disseminate and seed metastasis in distant organ. Our work identified Streptomycin (SM), a potent bactericidal antibiotic, as a new molecule capable of targeting non-adherent TIC from colon and breast cancer cell lines by inducing mitochondrial-dependent ferroptosis. SM-induced ferroptosis associates with profound alterations in mitochondrial morphology, such as swelling and cristae enlargement, coupled with hyperpolarization of mitochondrial membrane potential and production of mitochondrial ROS. The peculiar SM structure, and more particularly its aldehyde group, is essential for this mechanism. As such, the mere reduction of SM into dihydrostreptomycin abolishes its effect on TIC. This study reveals a new mechanism of action of SM that could help comprehend the molecular basis of TIC adaptation to inhospitable environments and pave the way for new treatment of advanced cancers.

## Introduction

Despite significant advances in cancer therapy, metastatic disease remains incurable in the vast majority of patients and make cancer the leading cause of death worldwide [1]. It is becoming increasingly clear that metastasis originates from a subset of disseminated cancer cells exhibiting stem-like properties [2] and marked similarities with tumor initiating cells (TIC). TIC phenotype is a plastic condition rather than a defined state, characterized by remarkable adaptation capabilities, self-renewal and multi-lineage differentiation capacity [3]. As such, they can disseminate and re-create a full-fledged tumor in unfavorable distant tissue environment [4]. For these reasons, targeting TIC became a major goal to eliminate cancer recurrence and improve clinical outcome of patients with advanced cancer. The limitations of this approach reside in the low susceptibility of TIC to cell death mediated by classical anticancer agents. Interestingly, salinomycin, an antimicrobial drug, selectively kills cancer stem-like cells by mediating ferroptosis, a non-apoptotic cell death process driven by reactive oxygen species (ROS) production and iron-dependent lipid peroxidation [5]. Along the same lines, antibiotics that induce mitochondrial dysfunction of TIC, also represent an appealing strategy for cancer treatment [6].

Cancer cells from either conventional cell lines or patient biopsies- can be grown at low cell density in non-adherent conditions and with depleted nutrients. In these unfavorable culture conditions, only a fraction of these cells can survive and form microtumor–like spheroids commonly named tumorspheres. In the case of solid tumor, this cell culture model is often used as a surrogate to evaluate tumorigenic potential [7]. It is well established that culture of cancer cells in suspension allows the enrichment of TIC whose resilience promotes their survival in unfavorable conditions [7]. We have previously shown that the Penicillin/Streptomycin (P/S) cocktail, commonly used in classical cell cultures, impinges on sphere forming ability [8]. Yet, P/S have no adverse effect on cell proliferation in monolayer cultures, suggesting a selected effect on cell subsets. Accordingly, P/S addition triggers a significant decrease in the number of Aldehyde Dehydrogenase (ALDH) positive cells. ALDH enzyme activity is widely used as a TIC marker in many solid tumors [9]. SM, a potent bactericidal antibiotic from aminoglycoside (AG) group, is generally administered for the treatment of individuals with moderate to severe infections. AG belong to a chemically diverse group of naturally secondary metabolites that bind to specific sites within the ribonucleoprotein complex in bacteria, where they interfere with protein synthesis and ultimately inhibit microbial growth [10]. SM structure stands out from other AG. The diaminoinositol unit is a streptidine, instead of 2-deoxystreptamin (2-DOS) or a streptamin, containing two highly basic guanidinium groups. Another molecular feature is the α-hydroxyaldehyde moiety that may create additional interactions with possible targets.

In this study, we identified SM as the sole AG capable of targeting non-adherent TIC by inducing ferroptotic cell death [11]. SM triggers significant mitochondrial dysfunction, ROS production, accumulation of the intracellular iron pool and lipid peroxidation, therefore recapitulating the hallmarks of ferroptosis. Remarkably, SM-resistant cells display a distinctive anti-oxidant gene expression signature that could represent a hallmark of highly adaptable TIC.

## Results

### SM inhibits TIC properties in suspension cultures

We have previously shown that the P/S cocktail inhibits sphere-forming ability in suspension cultures [8]. To identify which of these antibiotics was responsible for this effect, we tested them separately. We used three different human cancer cell lines: SW620 (liver metastasis from colon cancer), CRC1 (derived from patient with primary colon cancer tumor) [12] and MCF-7 (primary breast cancer). When grown in suspension, these cell lines form tumorspheres in seven to ten days (**figure 1.a**). By contrast with monolayer cultures, suspension cultures promote the enrichment of TIC, as illustrated by increased mRNA level of five genes involved in stemness (CD133, CD26, CD44, Oct4 and ALDH1A1) (**Figure S1.a**). As expected, SM decreased sphere forming ability in a dose dependent manner in all tested cell lines (**Figure 1.b**), while penicillin alone had no effect (**Figure S1.b**). SM was tested on multiple cancer cell lines, including CTC44, derived in the lab from circulating tumor cells isolated from patient with colorectal cancer [12], and three cell lines from different tissue origin (Gli4, glioblastoma; A549, lung cancer; Panc-1, pancreatic cancer). All tested cell lines displayed sensitivity to SM treatment in suspension culture, yet with variable sensitivity (**Figure 1.c, S1.c**). Gli4 and SW620 were the most sensitive cell lines (IC_50_ of 100 μM and 120 μM respectively), whereas Panc-1 was more resistant (lC_50_ around 300 μM). For CRC1 and MCF-7, the analysis revealed an IC_50_ to 165 μM and 180 μM. In agreement with our previous study, these observations suggest that SM exerts a pan cancer effect on sphere-forming cells, which mostly consists of TIC [13–15].

**Figure 1:**
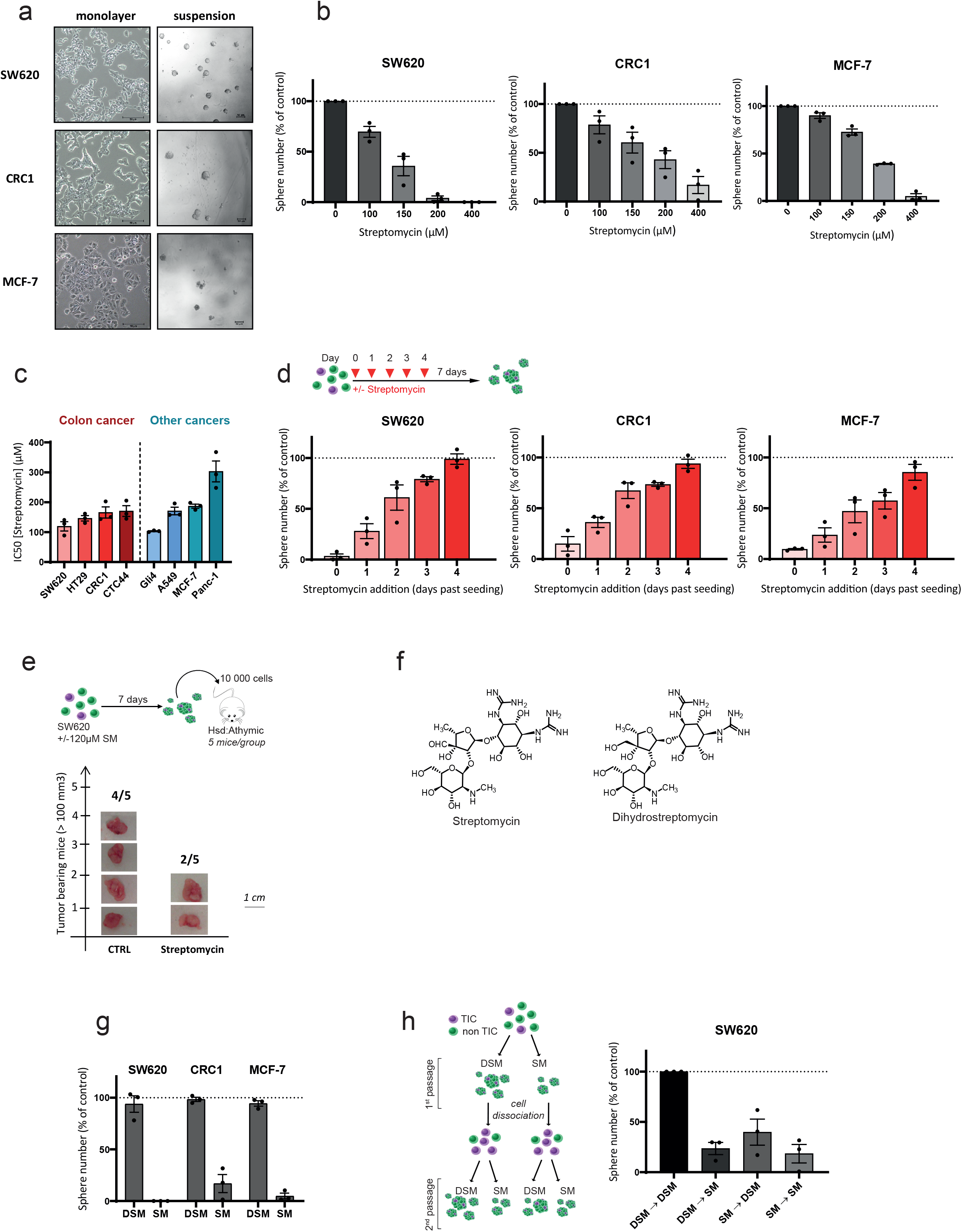
SM inhibits TIC properties in suspension culture. **a.** Representative pictures of SW620, CRC1 and MCF-7 cells cultured in monolayer and suspension. Scale bar 50 μM. **b.** Dose-response effects of SM on sphere forming ability on SW620, CRC1 and MCF-7 in suspension culture. Results are expressed in fold change compared to control and are mean ± SEM of three distinct experiments. **c.** IC_50_ of SM tested on different cell lines in suspension culture. Dose-response assays of SM were performed for each cell line and IC_50_ were calculated by nonlinear fitting parameters. Results are mean ± SEM of three distinct experiments. **d.** Impact of SM on sphere nucleation. SM at 200 μM (SW620) or 400 μM (CRC1, MCF-7) was added at different times following cell seeding (day 0, 1, 2, 3 or 4) in suspension culture. Sphere number was quantified at day 7. Results are expressed in fold change compared to control and are mean ± SEM of three distinct experiments. **e**. SM treatment decreases tumor initiation *in vivo.* The graph represents the number of tumor bearing mice for each group (five mice per group) following subcutaneous injection of SW620 cells pre-treated or not with SM at 120 μM. Scale bar 1 cm. **f.** chemical structure of SM and DSM. **g.** DSM does not impact sphere forming ability in SW620, CRC1 and MCF-7 cells. 400 μM of either SM or DSM were added in suspension culture and sphere forming ability was quantified. **h**. SM impairs self-renewal. SW620 cells were passaged once in suspension culture with either DSM or SM at 150 μM. Following 7 days, spheres were dissociated and cells were passaged a second time with DSM or SM. Following seven more days, the sphere number was evaluated for each condition. Results are expressed in fold change compared to control (DSM→DSM) and are mean ± SEM of three distinct experiments.

Next, we sought to understand whether SM impacts on sphere nucleation (i.e. the initiation of sphere formation) or on sphere growth. We treated SW620, CRC1 and MCF-7 cells with SM (200 μM for SW620, 400 μM for CRC1 and MCF-7) at different time points after cell seeding in suspension. Seven days after the beginning of the experiment, spheres were counted (**Figure 1.d**). In all tested cell lines, the maximal effect was obtained when SM was added at the time of cell seeding (day 0). When SM addition was delayed, its effect was gradually lost and SM adjunction became inefficient after day 4. Importantly, we did not notice any change of average sphere size amongst all tested conditions. This data suggests that SM impacts “sphere nucleation” -which relies on TIC phenotype– rather than “sphere growth”, fueled by more differentiated cells such as progenitors.

Next, we evaluated the impact of SM pre-treatment on *in vivo* tumor initiation potential, a distinctive feature of TIC. We inoculated immunodeficient mice (athymic nude) with suspension cultures of SW620 cells pre-treated or not with an IC_50_ concentration of SM (**Figure 1.e**). Thirty days later, tumor xenografts (diameter > 100mm^3^) were counted and harvested. Consistently with sphere formation assays, SM pre-treatment decreased by 50% tumor initiation potential.

Dihydrostreptomycin (DSM) is an antibiotic used in veterinary medicine that shares strong structure similarities with SM and nearly identical bactericidal properties [16]. DSM differs from SM by the reduction of the aldehyde group of the streptose moiety into a primary alcohol group (**Figure 1.f**). Interestingly, whereas this compound keeps a bacterial activity similar to SM, it did not display any effect on sphere forming-ability on SW620, CRC1 and MCF-7 cell lines at 400 μM (**Figure 1.g**). It is well known that aldehyde-containing drugs can form additional interactions with their target thanks to the formation of a reversible covalent bond with amino groups. This result suggests that the aldehyde group of SM could be involved in the mechanism of action against TIC. We then decided to use DSM as a negative control for all further experiments.

An important feature of TIC properties includes self-renewal ability. We have shown in SW620 cells that SM could interfere with this capacity (**Figure 1.h**). In this experiment, cells were passaged once in suspension culture with either DSM or SM at 150 μM. Upon seven days of culture, spheres (very few for SM condition) were dissociated and the same number of living cells were serially passaged with either DSM or SM. Following seven additional days of culture, the sphere number was evaluated for each condition. SM treatment in the second passage, decreases sphere number by 80% in comparison with DSM-treated control cultures. Interestingly, SM treatment followed by DSM treatment in the second passage did not rescue sphere formation, indicating that SM either alters self-renewal ability of TIC or kill most of them.

Importantly, both SM and DSM did not affect cell growth nor viability in monolayer culture (**Figure S1d & S1.e**). This substantiates the selectivity of SM over non-adherent TIC in cancer lines from different tissue origins.

### Sphere forming inhibition by SM is linked to its specific chemical structure

SM belongs to the AG class but stands out from other AG members as this compound is the only one having a streptidine genin instead of streptamine or 2-DOS (**Figure 2.a**). Nevertheless, we tested five other AG on their ability to inhibit sphere formation in suspension culture: neomycin, gentamicin, tobramycin, spectinomycin and paromomycin. These five antibiotics display either 2-DOS genin (neomycin, gentamicin, tobramycin and paromomycin) or streptamine genin (spectinomycin). In three tested cancer cell lines, these compounds had no effect on sphere formation potential as compared with SM at the same concentrations (**Figure 2.b**). Therefore, the distinctive structure of SM seems to be crucial for its effects on TIC, in particular its aldehyde group as emphasized above.

**Figure 2:**
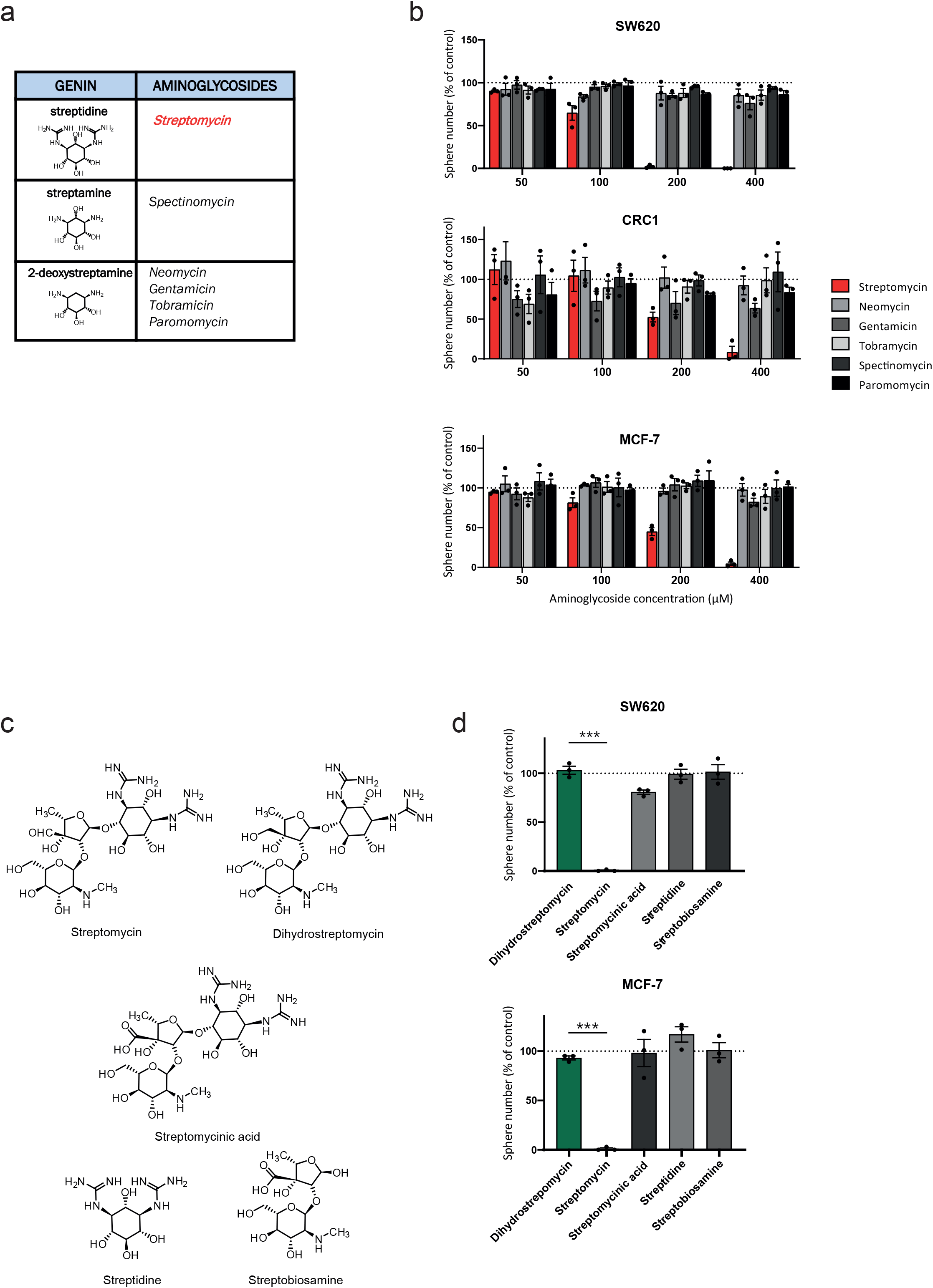
Sphere forming inhibition lies in SM structure uniqueness. **a**. SM displays atypical structure compared to other AG. **b.** Inhibition of sphere forming ability is not a common feature of AG. Dose response effect of several AG on sphere formation was evaluated in SW620, CRC1 and MCF-7 cells. Results are expressed in fold change compared to control (untreated cells) and are mean ± SEM of three distinct experiments. **c.** chemical structure of SM derivatives. **d.** Dose response effects of SM derivatives on sphere formation on SW620 and MCF-7 cell lines. Results are expressed in fold change compared to control. n=3 biological replicates. Mean ± SEM, ***p-value < 0.001. One-way Anova followed by multiple comparisons.

In order to better understand the importance of SM structure on sphere formation ability, we have synthesized three SM derivatives (**Figure 2.c** and **Figure s2**). The first one is streptomycinic acid, where the aldehyde group of the streptose moiety of SM has been oxidized into a carboxylic acid. This compound was prepared starting from commercially available streptomycin upon oxidation in the presence of bromine following a previously reported procedure [17]. Streptomycinic acid was obtained successfully in 73% yield. The two other derivatives represent the two parts of the SM structure: streptidine (the genin containing the guanidine group) and streptobiosamine (the association of the two sugar molecules, streptose and *N*-methyl-L-glucosamine). These two compounds were obtained upon hydrolysis of streptomycin in acidic conditions (H_2_SO_4_) in 33% and 27% yields, respectively. In SW620 and MCF-7 cell lines, the three synthesized compounds, streptomycinic acid, streptamine and streptobiosamine, did not affect suspension culture at 200 μM (SW620) or 400 μM (MCF-7) as opposed with native SM that abrogates sphere formation within this range of concentration (**Figure 2.d**).

Altogether, these results support the notion that the effect of SM on TIC is not a generic feature of AG but rather a specificity that lies in its peculiar chemical structure.

### SM-resistant cells deploy a transcriptional anti-oxidative stress response program

In order to give us some insights about the molecular mechanisms by which SM prevents sphere formation, we compared gene expression profile from IC_50_ SM-treated cells with DSM-treated cells by means of RNA-sequencing (RNA-Seq). SW620 cells were seeded in suspension conditions and total RNA was collected after two days of culture in presence of DSM 120 μM or SM 120 μM and compared to mock-treated cells (CTRL). Bioinformatic analysis did not reveal any change between the control cells and the DSM treated cells. However, we identified a SM-specific gene signature composed of eight genes that were upregulated. (**Figure 3.a & b).** Remarkably, all of these genes are involved in cell detoxification processes, suggesting they are involved in cell resistance to SM. The induction of these genes upon SM treatment was confirmed by quantitative Polymerase Chain Reaction (qPCR). Two sets of primers were used for each gene, in order to quantify the level of both mature and immature transcripts. qPCR analysis showed an overexpression of the target genes for both mRNA subtypes (**Figure 3.c**), implying that upregulation results from increased transcriptional activity rather from enhanced transcripts stability. Importantly, SM treatment did not upregulate these genes in monolayer culture (**Figure s3.a**). These results suggest that surviving TIC induce a cellular transcriptional program that prevents oxidative stress.

**Figure 3:**
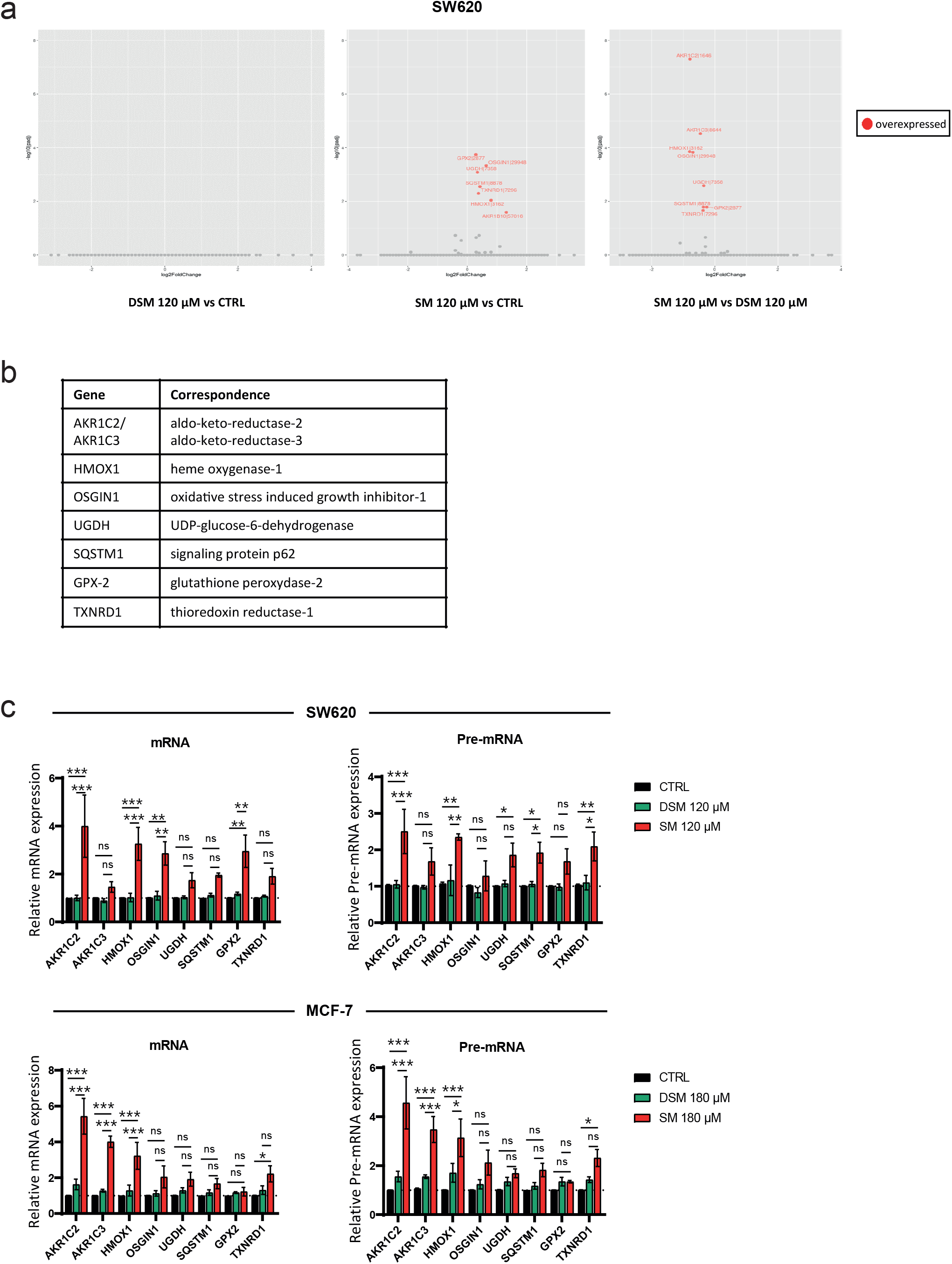
Surviving cells deploy a transcriptional anti-oxidative stress response program. **a**. DESeq2 analysis of RNA-Seq data, implemented with EdgeR. SW620 cells were cultured in suspension for 2 days before total RNA extraction and RNA sequencing. **b**. Table presenting the 8 upregulated genes in SM treated cells as compared with CTRL or DSM treated cells. **c**. validation of RNA-Seq by qPCR in SW620 and MCF-7 cells. mRNA level of genes was evaluated for mature and immature mRNA. Mean ± SEM of three biological replicates. ns not significant, *p-value < 0.05, **p-value < 0.01, ***p-value < 0.001. Two-way Anova followed by multiple comparisons.

Transcriptional analysis thus suggests a causal link between cell detoxification ability and resistance to SM. This observation is in line with the ability of AG to target mitochondria and produce reactive oxygen species [18].

### SM induces mitochondrial dysfunction

Recent reports suggests that antibiotics targeting prokaryotic ribosome could specifically impact mitochondrial biogenesis and activity in TIC and trigger cell death [19-21]. In order to test the impact of SM on mitochondrial biomass, we first employed MitoTracker Green (MTG) staining followed by flow cytometry analysis. The cells were seeded in suspension cultures and treated with either DSM or SM at their respective IC_50_ concentration (120 μM for SW620 and 180 μM for MCF-7). After six days (SW620) or ten days (MCF-7) of culture, MTG staining revealed no significant difference between SM-and DSM-treated cells (**Figure 4.a**). In addition, quantitative PCR analysis of mitochondrial DNA (mtDNA) copy number confirmed that total mitochondrial mass was not altered by SM (**Figure S4.a**).

**Figure 4:**
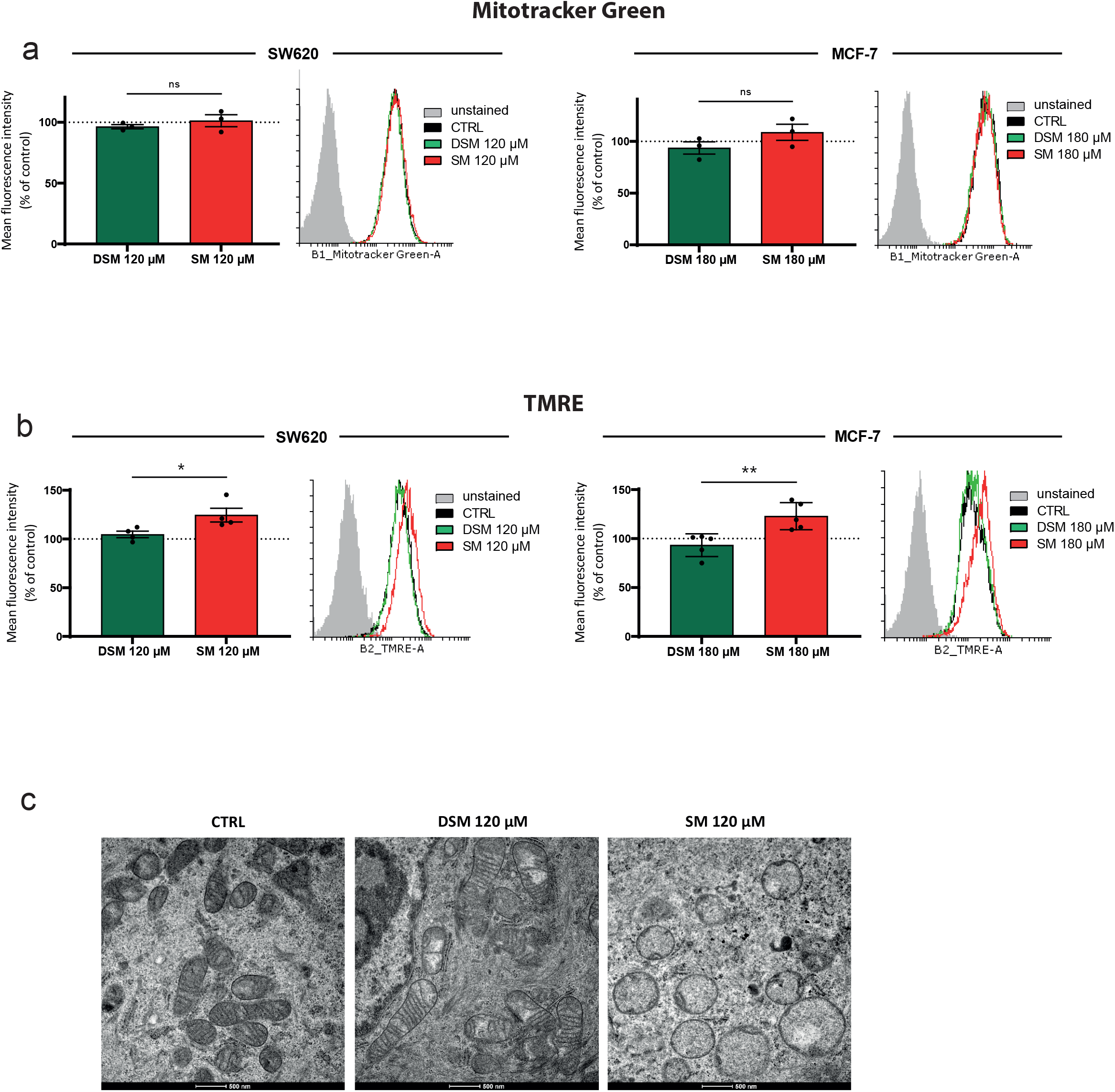
SM induces mitochondrial dysfunction. **a.** SM doesn’t impact mitochondrial biomass in suspension culture. Flow cytometry quantification of MTG (50 nM) mean fluorescence intensity in cells cultured in suspension with either DSM or SM at IC50 (120 μM for SW620 and 180 μM for MCF-7). Results are expressed in fold change compared to control. n=3 biological replicates. Mean ± SEM, ns not significant. Two-sided unpaired T-test. The histogram is a representative biological replicate (for each cell line). **b**. SM treatment induced hyperpolarization of MMP. Flow cytometry was performed to measure MMP by TMRE probe (20 nM) in SW620 and MCF-7 cells treated with DSM or SM in suspension culture. Results are expressed in fold change compared to control. n=3 biological replicates. Mean ± SEM, *p-value < 0.05, **p-value < 0.01. Two-sided unpaired T-test. The histogram illustrates one representative biological replicate. **c**. SM treatment induces mitochondrial morphological changes in suspension culture. Representative ultrastructure of mitochondria from SW620 treated with DSM or SM at 120 μM in suspension culture by TEM. Scale bar 500 nm.

Next, we monitored the effect of SM on mitochondrial membrane potential (MMP). We employed TMRE staining, a cell permeant, positively-charged dye that accumulates in active mitochondria. In SW620 and MCF-7 suspension culture, TMRE staining followed by flow cytometry showed a significant increase in MMP polarization in SM-treated cells as compared to DSM treated cells (**Figure 4.b**). We assessed mitochondrial morphology using transmission electron microscopy (TEM) in SW620 cells cultured in suspension with DSM or SM (120 μM) for six days. In SM treated cells, TEM revealed severe alterations in mitochondrial morphology with mitochondrial swelling and cristae vanishment (**Figure 4.c**). By contrast, SM treatment of monolayer culture at higher concentration (400 μM) did not promote any morphological difference (**Figure S4.b**).

Together, these results demonstrate that SM treatment affects mitochondrial function rather than mitochondrial biogenesis.

### The iron chelator deferoxamin inhibits the effect of SM on sphere formation

To decipher the mechanism by which SM triggers TIC death, we employed specific inhibitors targeting five major cell death types: apoptosis (Z-VAD-FMK), anoïkis (Y-27632), necroptosis (necrostatin-1, Nec-1), autophagy (chloroquine, CQ) and ferroptosis (deferoxamin, DFO). Their molecular mechanism of action is described in **Figure 5.a**. These inhibitors were used in three CRC cell lines (MCF-7, SW620, CRC1) at concentrations that are not toxic for cells in these culture conditions (**Figure S5.a**). The iron chelator DFO was the only compound that showed a significant dose-dependent rescue of sphere formation in SM-treated cells in the three tested cell lines (**Figure 5.b & c**). These findings strongly suggested that SM triggers TIC death through ferroptosis.

**Figure 5:**
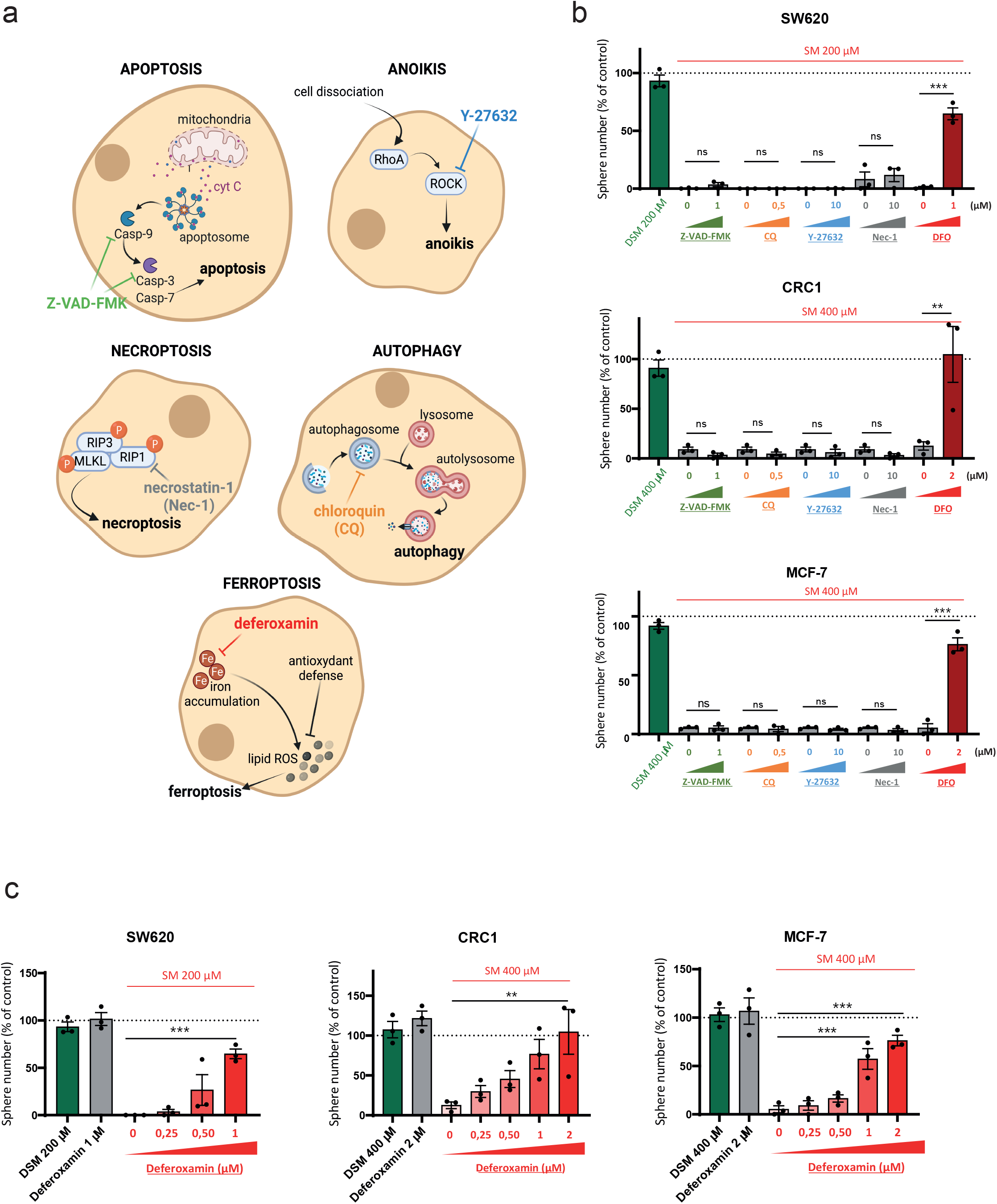
The iron chelator deferoxamin inhibits the effect of SM on sphere formation. **a**. Simplified illustration of the mechanism of action of different cell death inhibitors. Z-VAD-FMK is a cell-permeant pan caspase inhibitor that irreversibly binds to the catalytic site of caspase proteases and can inhibit induction of apoptosis. Y-27632 is a selective small molecule inhibitor of Rho-associated kinase (ROCK) involved in cell dissociation-induced anoikis. Necrostatin-1 (Nec-1) is a RIP1-targeted inhibitor of necroptosis. Chloroquine (CQ) prevents autophagosome formation during autophagy process. The iron chelator deferoxamin (DFO) is a ferroptosis inhibitor by preventing iron accumulation. **b**. DFO inhibits the effect of SM on sphere formation. Cell death inhibitors were used in combination with SM (200 μM for SW620, 400 μM for CRC1 and MCF-7) in suspension culture. Spheres were counted after 7 days. Results are expressed in fold change compared to control. n=3 biological replicates. Mean ± SEM, **p-value < 0.01, ***p-value < 0.001. One way Anova followed by multiple comparisons. **c**. DFO rescues sphere forming ability in dose-dependent manner. Cells in presence of SM (200 μM for SW620, 400 μM for CRC1 and MCF-7) were co-treated with increasing concentrations of DFO. Sphere number was evaluated after 7 days. Results are expressed in fold change compared to control. n=3 biological replicates. Mean ± SEM, **p-value < 0.01, ***p-value < 0.001. One way Anova followed by multiple comparisons.

### SM kills TIC through mitochondrial dependent ferroptosis

Considering the substantial impact of SM on mitochondrial morphology, we wondered whether ferroptosis induction could originate from this organelle. Indeed, mitochondria plays a central role in ROS production as well as in iron metabolism [22], and their implication in ferroptosis has been previously documented [23]. First, we evaluated the impact of SM on mitochondrial ROS production in SW620 and MCF-7 cells seeded in suspension. Staining of both cell lines with MitoSOX Red, a dye commonly used to quantitate mitochondrial superoxide, showed an important increase in mitochondrial ROS production (**Figure 6.a**). Global ROS production was also investigated by CM-H_2_DCFDA staining. This dye becomes fluorescent after cellular oxidation and removal of acetate groups by cellular esterases. Our results revealed an increase of H_2_DCFDA staining after SM treatment in SW620 (**Figure S6.a**). This suggests that ROS production originates mainly from mitochondria. Consistently, N-acetylcysteine (Nac) and Mito-Tempo (MT), respectively a general antioxidant and a mitochondrial targeted antioxidant, were both able to counteract SM-mediated redox imbalance and restore sphere formation potential (**Figures 6.b, S6.b**). Of note, MT was toxic in suspension culture for MCF-7 cells (even at low concentration). Therefore, we modified our protocol accordingly: cells were pretreated with MT (100 μM) in monolayer culture before passaging in suspension cultures with a range of SM concentrations. Tempol (4-hydroxy-TEMPO), a superoxide dismutase mimetic, restored either partially (CRC1, MCF-7) or totally (SW620) sphere formation ability in SM-treated suspension cultures (**Figure S6.c**). These results indicate that ROS production is an important event in SM-induced TIC death mechanism.

**Figure 6:**
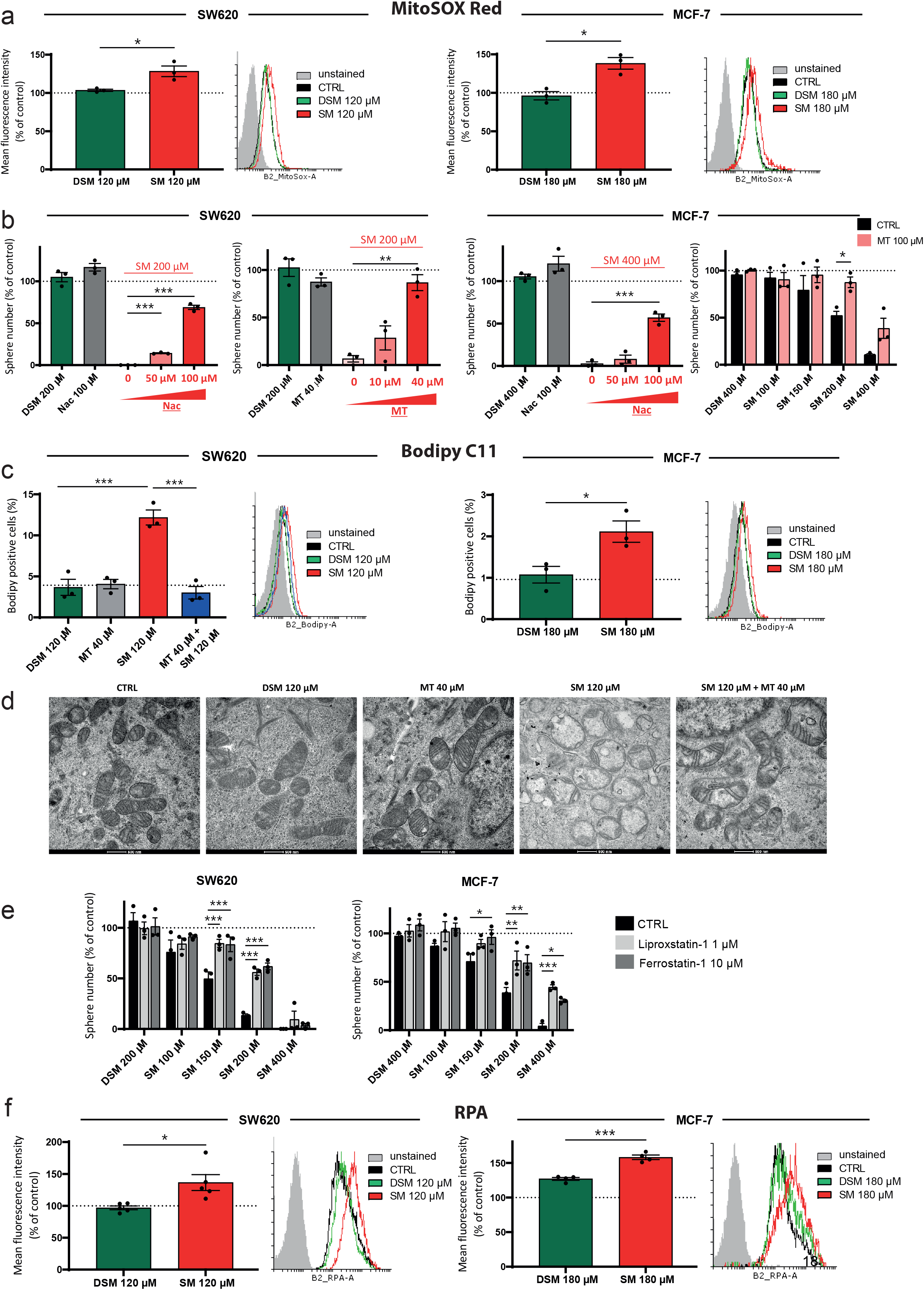
SM kills TIC through mitochondrial dependent ferroptosis. **a.** Flow cytometry quantification of MitoSOX Red (5 μM) mean fluorescence intensity in cells cultured in suspension condition with DSM and SM at 120 μM (SW620) or 180 μM (MCF-7). Results are expressed in fold change compared to control. n=3 biological replicates. Mean ± SEM, *p-value < 0.05. Two-sided unpaired T-test. Histogram representative of triplicates. b. ROS inhibitors restore sphere formation ability. Cells treated with SM at 200 μM (SW620) or 400 μM (MCF-7) in suspension condition were co-treated with increasing concentrations of either N-acetylcysteine (Nac) or Mito-Tempo (MT). Results are expressed in fold change compared to control. n=3 biological replicates. Mean ± SEM, **p-value < 0.01, ***p-value < 0.001. One way Anova followed by multiple comparisons. Right: MCF-7 were pretreated with MT at 100 μM for 3 days, then passed in suspension culture with increasing concentrations of SM. Mean ± SEM, *p-value < 0.05. Two way ANOVA followed by multiple comparisons. c. SM treatment promotes lipid peroxidation in suspension culture. Flow cytometry quantification of Bodipy-C11 (10 nM) positive cells in cells cultured in suspension condition with DSM and SM at 120 μM (SW620) or 180 μM (MCF-7). Left: SW620 cells were co-treated with MT at 40 μM. Results are expressed in fold change compared to control. n=3 biological replicates. Mean ± SEM, *p-value < 0.05, ***p-value < 0.001. SW620: One way Anova followed by multiple comparisons. MCF-7: Two-sided unpaired T-test. d. The mitochondrial antioxidant MT rescues normal mitochondrial morphology in suspension culture. Representative ultrastructure of mitochondria from SW620 suspension cultures treated with DSM or SM at 120 μM with or without addition of MT (40 μM). Scale bar 500 nm. e. Lipid ROS inhibitors counteract SM effect on sphere formation. SW620 and MCF-7 cell lines were pretreated in monolayer with Lip-1 (1 μM) or Fer-1 (10 μM). After 3 days, cells were passaged in suspension culture with increasing concentrations of SM. Results are expressed in fold change compared to control. n=3 biological replicates. Mean ± SEM, *p-value < 0.05, **p-value < 0.01, ***p-value < 0.001. Two way Anova followed by multiple comparisons. f. Mitochondrial iron pool is increased after SM treatment in suspension culture. Flow cytometry quantification of RPA (1 μM) mean fluorescence intensity in cells cultured in suspension with IC50 concentrations of either DSM or SM (120 μM for SW620 or 180 μM for MCF-7). Results are expressed in fold change compared to control. n=3 biological replicates. Mean ± SEM, *p-value < 0.05, ***p-value < 0.001. Two-sided unpaired T-test. Histogram is representative of three independent experiments.

Another important hallmark of ferroptosis is lipid peroxidation, often considered as the final event in this process. We evaluated the levels of lipid peroxidation in SW620 and MCF-7 cells by flow cytometry upon incubation with the fluorescent dye BODIPY C11. SM treatment almost doubled the number of BODIPY C11 positive cells, confirming the significant increase of lipid peroxidation (**Figure 6.c**). In SW620 cells, co-treatment with the mitochondrial antioxidant MT restored lipid peroxidation to the basal state, demonstrating the importance of mitochondrial metabolism in this process. The mitochondrial origin of SM-induced ferroptosis in SW620 cells was also confirmed by TEM observations showing significant restoration of mitochondrial morphology when these cells were cultured in suspension in presence of MT (**Figure 6.d**).

Lipid peroxidation and ferroptosis can be prevented by liproxstatin-1 (Lip-1) or ferrostatin-1 (Fer-1). As these molecules were toxic for suspension cells, cancer cell lines (SW620, CRC1 and MCF-7) were pretreated in monolayer culture with 1 μM of Lip-1 or 10 μM of Fer-1 before passaging them in suspension in growing concentrations of SM. In all SM-treated conditions, pre-treatment with both inhibitors restored significantly sphere formation (**Figures 6.e, S6d**), emphasizing the importance of lipid peroxidation in SM-mediated cell death.

Another crucial driver of ferroptosis is intracellular iron accumulation. Iron induces the generation of hydroxyl and peroxyl radicals through the Fenton reaction, which promotes lipid ROS production. Calcein-AM and RPA dyes were used to monitor cytoplasmic and mitochondrial free iron pools, respectively. SM treatment of either SW620 or MCF-7 cells resulted in a significant increase in free iron pools in both cellular compartments (**Figures 6.f, S6.e**).

Finally, we evaluated the role of GPX-4, a major antioxidant enzyme that regulates the cellular redox potential by using GSH to detoxify lipid peroxidation. Dysregulation of GPX-4 in ferroptosis has been largely documented [24]. However, GPX-4 does not seem implicated in SM-mediated ferroptosis as revealed by our analysis showing no effect on the GSH/GSSG ratio in SW620 cells (**Figure S6.f**).

Altogether, these results demonstrate that SM triggers unconventional ferroptosis of TIC in suspension cultures through a mechanism leading to dysregulation of iron metabolism.

### Discussion

In this study, we show that SM targets non-adherent TIC and promotes ferroptosis. We unraveled a unique transcriptional program that likely contributes to cell detoxification and SM resistance. We demonstrate the importance of SM structural features in triggering this mechanism of cell death, in particular the aldehyde group. Chemical reduction of the former in DSM-provided us an ideal control for *in vitro* and *in vivo* experiments. Finally, we characterized SM-induced ferroptosis and showed that changes in iron metabolism in SM-treated cells lead to increased mitochondrial ROS production.

The main features characterizing ferroptosis – such as increased lipid peroxidation-make cancer cells more sensitive than normal cells to this cell death mechanism. TIC in particular may represent ideal targets due to their changes in lipid metabolism that allow them to adapt to nutrient-poor environments [25]. Iron dependence is another hallmark of TIC [5, 26, 27], which may also contribute to their sensitivity to SM-induced ferroptosis. Two other antibiotics, salinomycin and its derivative ironomycin, have been previously identified as ferroptotic inducers in TIC [5]. Yet, the underlying molecular mechanisms may differ from that triggered by SM. SM-induced mitochondria dysfunctions, which result in significant morphological changes and elevated mitochondrial oxidative stress, takes a central place in this process.

The relevance of mitochondrial dysfunction in ferroptosis is still under debate. Indeed, whereas Gaschler and colleagues showed that ferroptosis-sensitive HT-1080 fibrosarcoma cells without mitochondria are still responsive to ferroptotic signals [28], another study showed in the same cells that inhibition of mitochondrial electron transport chain (ETC) could rescue MMP and lipid peroxidation [29]. In most cases, ferroptotic cells display reduced mitochondrial size with disappearance of cristae and increased MMP [29]. In contrast, SM-treated cells exhibited swollen mitochondria, though with membrane hyperpolarization and decreased cristae number. This phenotype has already been found in a recent study that identified mitochondrial protein frataxin (FXN) as a regulator of ferroptosis in HT-1080 cells [30]. Thus, the relevance of mitochondria in the ferroptotic process may vary upon cellular context, in particular for cell types that rely on mitochondrial mechanism such as TIC [31].

Our results raise an important question regarding the mechanism by which SM triggers mitochondrial dysfunction. Our data showing that DSM and other aminoglycosides do not trigger TIC death rules out the involvement of mitochondrial translation inhibition. Furthermore, targeting mitochondrial ribosomes would lead to decreased mitochondria biomass and affect monolayer cell growth at high SM concentration. Yet, none of these defects was observed, suggesting that SM induced an alteration of mitochondrial function rather than biogenesis. The increased MMP that we identified with TMRE staining may originate from the alteration of voltage-dependent anion channels (VDAC). VDAC control the trafficking of ions and other metabolites between mitochondria and cytosol [32] and alteration of their activity can lead to ferroptosis [33]. Several studies suggest that SM and other AG are able to block stretch-activated channels. In particular, SM can prevent entry of calcium in cells by blocking transient receptor potential channels like TRPV1 in neurosensitive neurons [34] or TRPM7 in pulmonary fibroblasts [35]. While this has yet to be demonstrated, this type of channels represent potential SM targets, which inhibition could lead to severe functional and morphological changes in mitochondria.

In the course of this study, we uncovered a new transcriptional program that likely counteracts oxidative stress and allows the survival of a subpopulation of SM-treated TIC. Remarkably, this inducible program was restricted to eight genes that are involved in cell detoxification and/or ferroptosis inhibition. For instance, AKR1C2 and AKR1C3 are aldo-keto-reductases involved in NADPH-dependent reductions. HMOX1 has been associated with systemic iron homeostasis, stress response [36], and more recently with ferroptosis [37]. Surprisingly, GPX-4 that is tightly associated with ferroptotic process [24], was not identified among this limited set of genes. However, a member of the same family, GPX-2 (glutathione peroxidase-2), was found to be upregulated upon SM treatment. Finally, a recent study suggests that TXNRD1 (thioredoxin reductase-1) could act as a ferroptotic modulator in chronic myeloid leukemia [38]. Most certainly, enhancing expression of this specific set of genes requires a very selective transcriptional activator such as, for example, ZNF410, the NuRD component CHD4 [39]. Our findings pave the way for future studies aiming at identifying this factor for the design of new therapeutic strategies targeting oxidative stress resistant cells such as TIC.

Most importantly, our study reveals a new mechanism of action of SM that could be exploited in cancer biology. Since SM is already an FDA-approved drug used in clinic, it represents an ideal candidate for drug repositioning in anti-TIC therapy. Furthermore, SM has a pan-cancer activity, which reinforce its value in clinic. While side effects, such as ototoxicity, can be put forward to limit its use, preventing tumor spreading in advanced (e.g. stage 3 colorectal cancer) cancer patient remains a matter of life and death. Further, combined with chemotherapy, SM could also be of use to prevent bacterial infections in neutropenic patients.

## Acknowledgements

This work was generously supported by INCa (PLBio 2017-160), Ligue contre le Cancer, SIRIC Montpellier Cancer (INCa-DGOS-Inserm 6045), Fondation ARC and Cancéropole GSO. We thank Montpellier Genomix (http://www.mgx.cnrs.fr) sequencing facility, in particular Hugues Parinello and Xavier Mialhe, as well as iExplore animal facility, in particular Amelyne David and Oceane Gentilini.

We would like to thank Jean-Emmanuel Sarry (CRCT, Toulouse) and Laurent Le Cam (IRCM, Montpellier) for their helpful insights.

The authors declare no conflict of interest.

## Author contributions

A.D., M.D., F.M., H.G. designed experiments and analysed the results. H.G., S.R., L.B., B.Z., F.B., A.dG., C.C., C.B., M.B., S.H., A.C., C.P. A.B. performed experiments. A.D., M.D., F.M., H.G., X.M., J.P. performed data analysis. A.D., M.D., F.M., H.G. wrote the manuscript. All the authors reviewed the final version of the manuscript.

## Methods

### Cell lines

Patient-derived cell culture of colon cancer cells CRC1 were obtained from primary tumor of colorectal biopsies provided by CHU-Carémeau (Nîmes, France) within an approval protocol (ethical agreement no 2011-a01141-40). Signed informed consents were obtained from patients prior to samples acquisition in accordance with all ethical and legal aspects. CTC44 are circulating tumor cells derived from blood of metastatic chemotherapy-naïve stage IV colorectal patients. SW620, HT29, Gli4, MCF-7, Panc-1 and A549 cells were purchased from American Type Culture Collection (ATCC).

### Cell culture in monolayer

Cells were maintained at 37°C under humidified 5% CO2 in DMEM medium (Gibco) supplemented with 10% FCS (Eurobio) and 2 mM glutamine.

### Cell culture in suspension and sphere formation assay

Cells were plated at 100 cell/100μL in M11 medium (DMEMF12 (1:1) GlutaMAX medium, N2 Supplement, Glucose 0.3%, insulin 20 μg/mL, hBasic-FGF 10 ng/mL, hEGF 20 ng/mL) in low attachement plates treated with Poly-HEME at 10 mg/mL. For further experiments, spheres were incubated for 40 min at 37°C with Accumax solution (Sigma). For sphere formation assay, spheres > diameter were counted after 7-10 days following different treatments.

### Proliferation assay

Cell proliferation was monitored in real-time using the xCELLigence system E-Plate. Two thousands cells were seeded in wells with DSM or SM at 400 μM in DMEM medium. The impedance value of each well was monitored by the xCELLigence system for duration of 72 h and expressed as a cell index value.

### Cytotoxicity assay

A thousand of cell were seed in 96-well plate with decreasing doses of DSM or SM (1/3 dilution from 2 mM). After 96 h, cells were fixed for at least 1 h at 4°C in 10% TCA. After two washes with Milli-Q water, cells were incubated in 0.4% Sulforhodamine B (SRB) solution for 20 min at room temperature followed by three washes with acetic acid 1% solution. Absorbance was measured at 560 nm after resuspension in 10 mM Tris-Base.

### Tumor initiation assay

SW620 cells were grown in suspension with or without 120 μM SM for 7 days. Spheres were dissociated and 10 thousands living cells were subcutaneously injected into nude mice (Hsd:Athymic Nude-Foxn1nu nu/nu, 6 weeks, females, 5 mice per group) in Matrigel - DMEM (v : v). After 50 days, the mice were sacrificed and tumors were taken out. The number of mice bearing growing tumor (size > 100 mm^3^) was counted.

### RNA extraction and quantitate real-time PCR (RT-qPCR)

Total RNA was extracted using TRIzol reagent (Invitrogen) according to the manufacturer instructions. For RT-qPCR analyses, 1 μg of RNA was reverse- transcribed into cDNA using random hexamers (Invitrogen) and 1 U of MML-V reverse transcriptase (Invitrogen). Quantitative gene expression was performed using SYBR Green master mix (Roche) on LightCycler 480 Instrument (Roche). Results were normalized using the ΔΔCt method. Primer sequences are provided as follows: GAPDH-F: 5’-CCCACTCCTCCACCTTTGAC-3’ ; GAPDH-R: 5’-CCACCACCCTGTTGCTGTAG-3’; Actin-F: 5’- AGCACGGCATCGTCACCAACT-3’; Actin-R: 5’-TGGCTGGGGTGTTGAAGGTCT-3’; Oct4-F: 5’- GTGGAGAGCAACTCCGATG-3’; Oct4-R: 5’-TGCTCCAGCTTCTCCTTCTC-3’; CD44-F: 5’-TGGCACCCGCTATGTCGAG-3’; CD44-R: 5’-GTAGCAGGGATTCTGTCTG-3’; CD133-F: 5’-TGGGGCTGCTGTTTATTATTCT-3’; CD133-R: 5’-TGCCACAAAACCATAGAAGATG-3’; CD26-F: 5’-CAAATTGAAGCAGCCAGACA-3’; CD26-R: 5’-CACACTTGAACACGCCACTT-3’.

### Determining mitochondrial DNA (mtDNA copy number)

Quantification of the mtDNA copy number relative to the nuclear DNA (ncDNA) was realized using real-time PCR as previously described [40]. Real-time quantitative PCR to quantify mtDNA and ncDNA was performed on 10 ng nucleic acids, directly on total RNA isolated by TRIzol, by using the SYBR Green master mix (Roche) and the LightCycler 480 instrument (Roche), with the following program: 10 min at 95°C, 50 cycles of 10 s at 95°C, 15 s at 67°C and 15 s at 72°C. ncDNA was quantified by using the following primers: β2-microglobulin-F: 5’-TGCTGTCTCCATGTTTGATGTATCT-3’; β2-microglobulin-R: 5’-TCTCTGCTCCCCACCTCTAAGT-3’. Total mtDNA was quantified by amplifying a DNA domain within the D-loop of mtDNA by using the following primers: Universal-F: 5’-TTAACTCCACCATTAGCACC-3; Universal-R: 5’-GAGGATGGTGGTCAAGGGA-3.

### Measurement of mitochondrial biomass

Mitochondrial mass was measured by MitoTracker Green FM (MTG). After treatment with DSM and SM (120 μM for SW620, 180 μM for MCF-7), cells were cultured in suspension condition for 6 days (SW620) or 10 days (MCF-7) and spheres were dissociated with Accumax. Fifty thousand cells were stained with 50 nM MTG for 10 min at 37°C and the fluorescence emission was detected by flow cytometry.

### Measurement of mitochondrial membrane potential (MMP)

Level of MMP was measured with tetramethylrhodamine methyl ester (TMRE) probe. Cells were treated in suspension with SM or DSM (120 μM for SW620, 180 μM for MCF-7). After 6 days (SW620) or 10 days (MCF-7), spheres were dissociated and fifty thousand cells were stained with 20 nM TMRE at 37°C for 15 min. Excess TMRE was removed by washing the cells once with PBS/BSA 1% and analysis was done following flow cytometry.

### Transmission electron microscopy

Tissues were immersed in a solution of 2.5% glutaraldehyde in PHEM buffer (1X, pH 7.4) overnight at 4°C. They were then rinsed in PHEM buffer and post-fixed in a 0.5% osmic acid + 0.8% potassium Hexacyanoferrate trihydrate for 2 h at dark and room temperature. After two rinses in PHEM buffer, the cells were dehydrated in a graded series of ethanol solutions (30-100%). The tissues were embedded in EmBed 812 using an Automated Microwave Tissue Processor for Electronic Microscopy, Leica EM AMW. Thin sections (70 nm; Leica-Reichert Ultracut E) were collected at different levels of each block. These sections were counterstained with uranyl acetate 1.5% in 70% Ethanol and lead citrate and observed using a Tecnai F20 transmission electron microscope at 120 KV in the Institut des Neurosciences de Montpellier: Electronic Microscopy facilities, INSERM U 1298, Université Montpellier, Montpellier France.

### Determination of ROS production

Cells were seeded in suspension culture with SM or DSM at 120 μM (SW620) or 180 μM (MCF-7) for 6 days (SW620) or 10 days (MCF-7). Total ROS generation was measured with CM-H_2_DCFDA (Invitrogen) and mitochondrial ROS were assessed with MitoSOX Red (Invitrogen). Spheres were dissociated with Accumax and fifty thousand cells were incubated with 50 nM (SW620) or 500 nM (MCF-7) CM-H2DCFDA or 5 μM MitoSOX at 37°C for 10 min. Cells were harvested and analysis was done following flow cytometry.

### Assessment of lipid peroxidation using C11-BODIPY

Lipid peroxidation was measured with C11-BODIPY (581/591). Cells were seeded in suspension culture with SM or DSM at 120 μM (SW620) or 180 μM (MCF-7). After 6 days (SW620) or 10 days (MCF-7), spheres were dissociated and fifty thousand cells were incubated with 10 nM C11-BODIPY (581/591) at 37°C for 10 min. Cells were harvested and resuspended for flow cytometry.

### Determination of cytoplasmic and mitochondrial iron pool

The cytoplasmic chelatable iron pool was assessed using Calcein-AM and the mitochondrial iron pool was detected using Rhodamine B-[(1,10-phenanthroline-5-yl)-aminocarbonyl]benzyl ester (RPA), both specific Fe^2+^ specific fluorescent sensors. After treatment of cells with DSM or SM (120 μM for SW620, 180 μM for MCF-7) during 6 days (SW620) or 10 days (MCF-7) following sphere dissociation with Accumax, fifty thousand cells were harvested and incubated with 10 nM Calcein-AM or 1 μM RPA for 15 min at 37°C. Following two washes with PBS/BSA 1%, cytoplasmic and mitochondrial iron pool were measured by flow cytometry.

### GSH/GSSG ratio detection

SW620 cells were seeded in suspension culture for 6 days with DSM or SM at 120 μM. After sphere dissociation with Accumax, five thousand cells were plated in white 96-well plate. GSH and GSSG were quantified using the GSH/GSSG-Glo Assay kit (Promega), according to the manufacturer’s protocol. Luminescence in relative light unit (RLU) was measured with a plate reader (Tecan Infinite M200) and the following formula was used to calculate GSH/GSSG ratio: (Net treated total glutathione RLU – Net treated GSSG RLU) / (Net treated GSSG RLU / 2).

### RNA-Sequencing

Total RNA from SW620 cells cultured in suspension in 3 biological replicates (Control, DSM 120 μM, SM 120 μM) were extracted using TRIzol Reagent (Invitrogen) following manufacturer procedure. Quality and quantity were evaluated by electrophoretic migration and spectrometry. 200 ng total RNA per sample was sequenced by the platform MGX. Library was prepared using Universal Plus mRNA-seq by NuGEN. Validation of the library was performed on Fragment Analyzer (Standard Sensitivity NGS kit) and using qPCR (ROCHE Light Cycler480, France). Total RNAs were sequenced on NovaSeq 6000 from Illumina. Biological validation of RNA-Seq results was performed using quantitative Polymerase Chain Reaction (qPCR) (primer sequences in **sup Table 1**).

## SUPPLEMENTAL INFORMATION for

**Figure S1.**
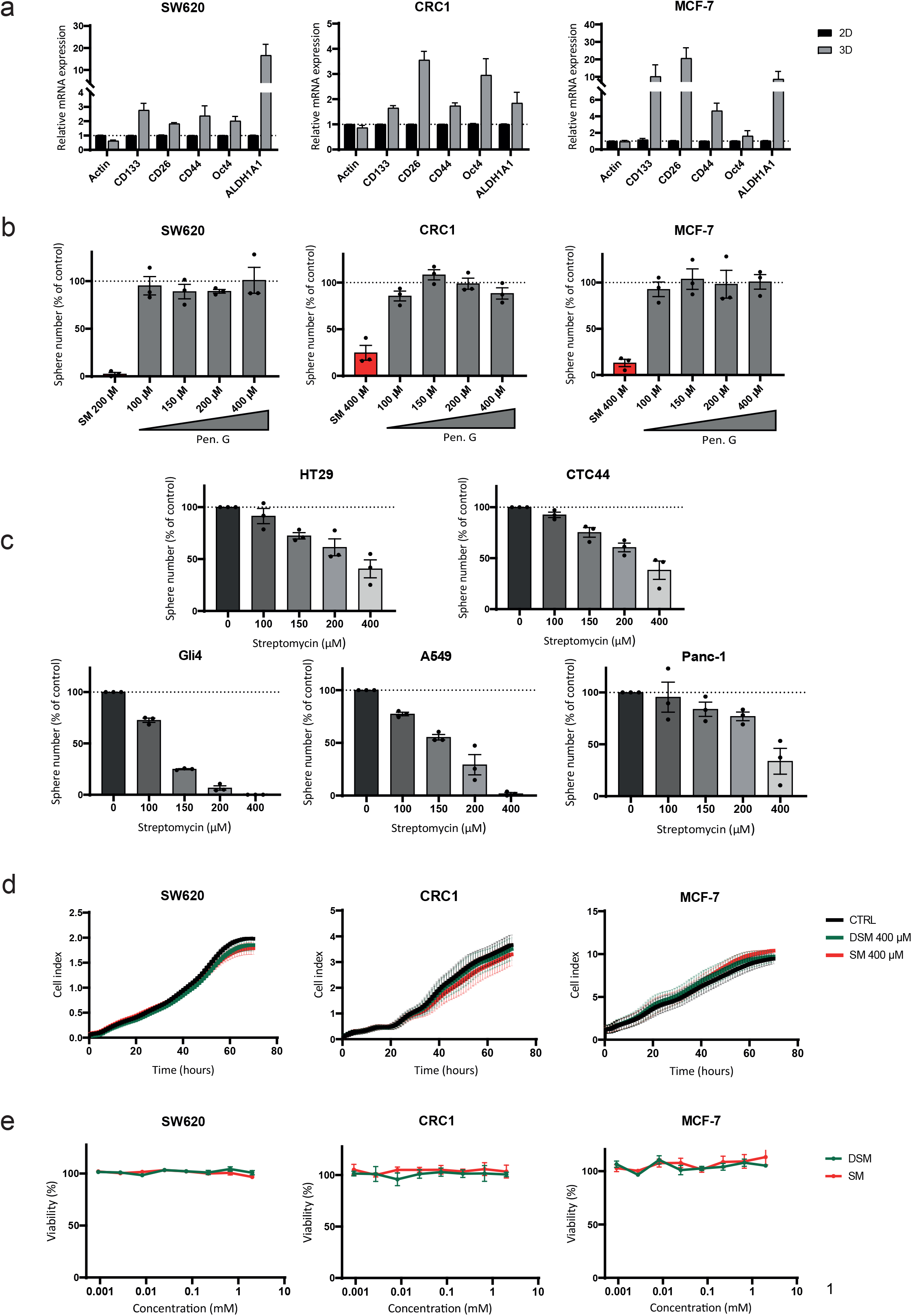
**a**. TIC markers are enriched in suspension culture. mRNA level of genes involved in stemness was evaluated by RT-qPCR in SW620, CRC1 and MCF-7 cells cultures in monolayer or suspension conditions in absence of antibiotic. Mean ± SEM of three biological replicates. **b**. Penicillin G does not impact sphere forming ability. Increasing concentrations of Penicillin G were added in suspension cultures of SW620, CRC1 and MCF-7. Spheres were counted following 7 days. Results are expressed in fold change compared to control and are mean ± SEM of three distinct experiments. **c.** Dose-response effects of SM on sphere forming ability on HT29, CTC44, Gli4, A549 and Panc-1 in suspension culture. Results are expressed in fold change compared to control and are mean ± SEM of three distinct experiments. **d.** DSM and SM do not impact cell growth in monolayer culture. Cell proliferation of SW620, CRC1 and MCF-7 was evaluated in the presence or absence of antibiotics (SM or DSM at 400 μM). Results (expressed in cell index) are the mean ± SEM of a representative experiment (out of three separate experiments). **e**. DSM and SM are not toxic in monolayer culture. DSM and SM toxicity were evaluated on SW620, CRC1 and MCF-7 cells. Results are expressed in fold change compared to control (untreated cells) and are mean ± SEM of three distinct experiments.

**Figure S2.**
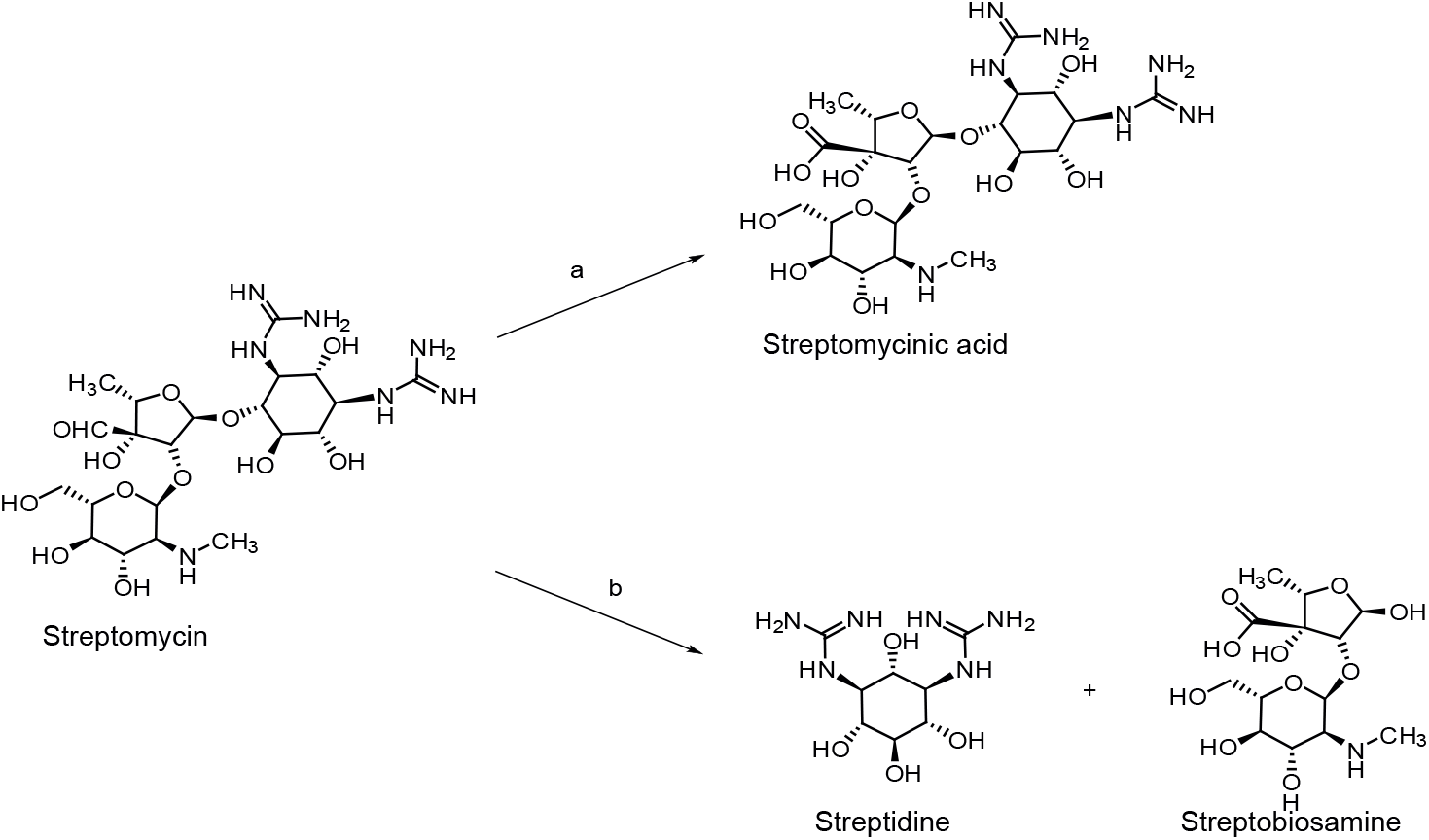
Synthesis of streptomycinic acid, streptidine and streptobiosamine starting from streptomycin. Reagents: a) Br_2_, H_2_O, 5 days, r.t.; b) H_2_SO_4_, H_2_O, 48h, 37°C.

**Figure S3.**
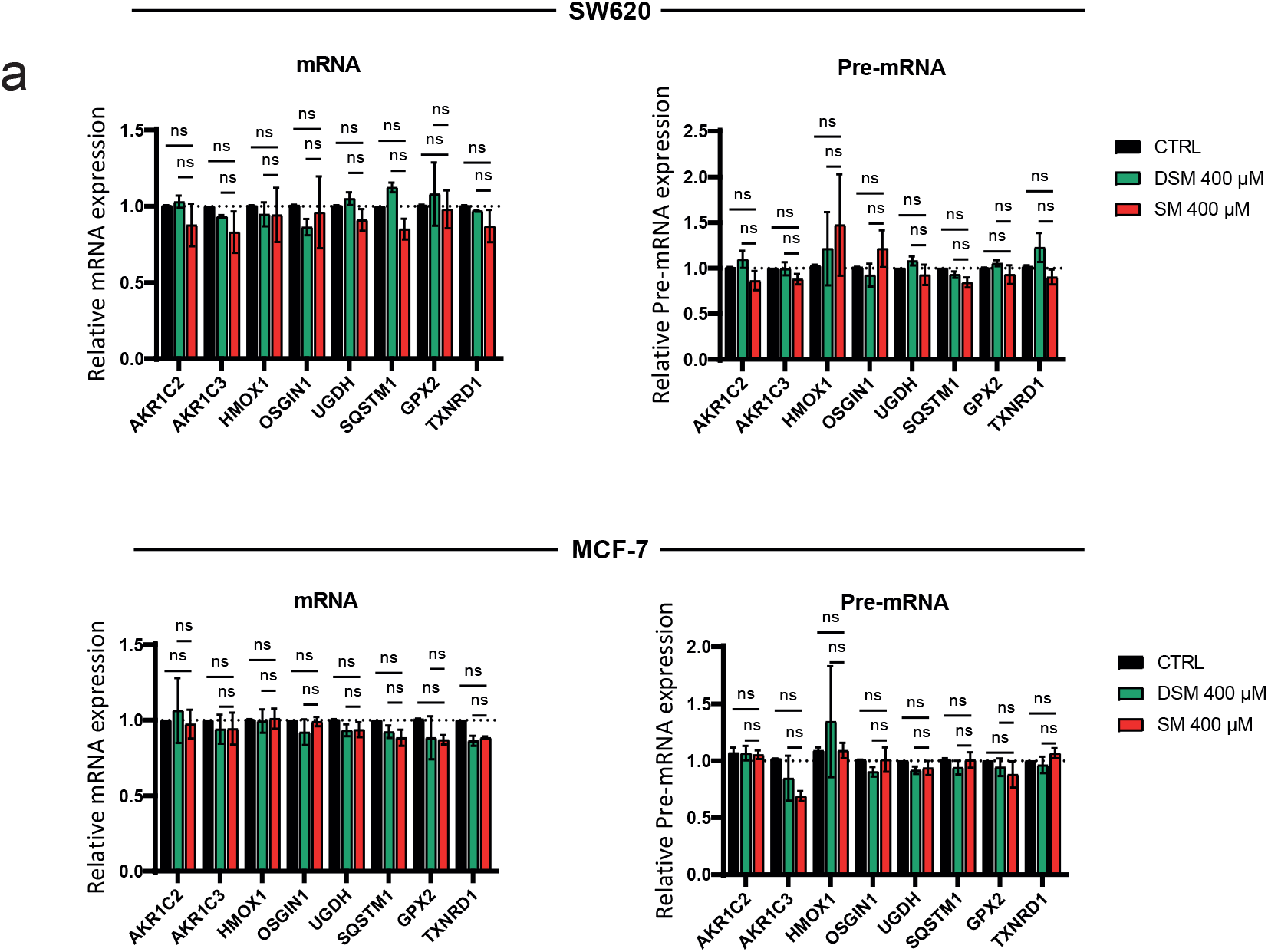
**a**. Gene signature is absent in SM-treated monolayer culture. SW620 and MCF-7 cells were treated with either SM or DSM at 400 μM in monolayer culture for 4 days. mRNA level was evaluated for mature and immature (unspliced) mRNA. Mean ± SEM of three biological replicates. ns not significant. Two-way Anova followed by multiple comparisons.

**Figure S4.**
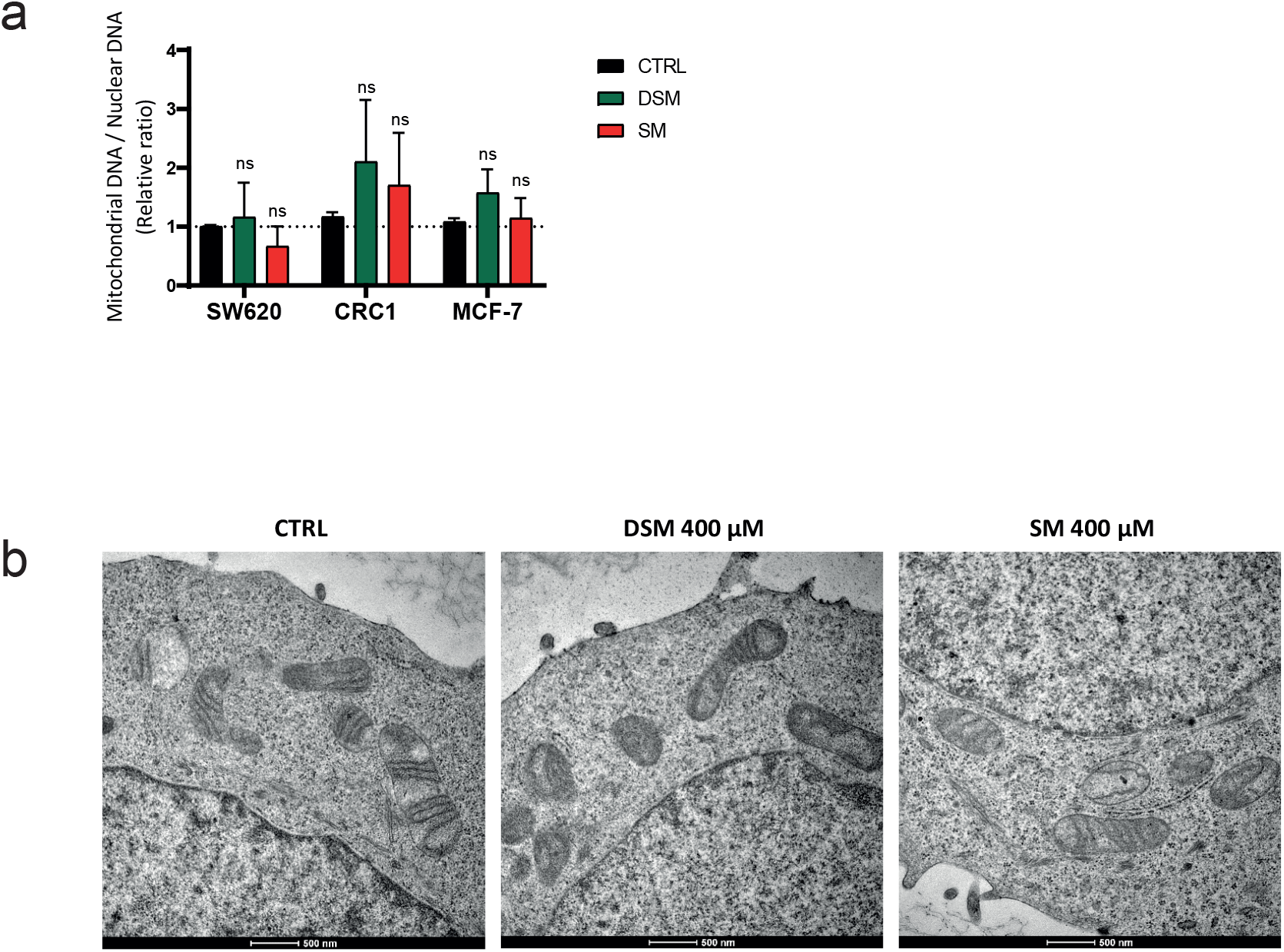
**a**. mtDNA copy number is not impacted by SM in suspension culture. mtDNA/nucDNA ratio in cells treated with SM and DSM in suspension culture (120 μM SW620, 165 μM CRC1, 180 μM MCF-7). PCR quantification was carried out following RNA extraction. n=3 biological replicates. Mean ± SEM, ns not significant. One way Anova followed by multiple comparisons. **b**. SM treatment does not impact mitochondria morphology in monolayer culture. Ultrastructure of mitochondria from either SM- or DSM-treated monolayer culture of SW620 (400 μM) by TEM. Scale bar 500 nm.

**Figure S5.**
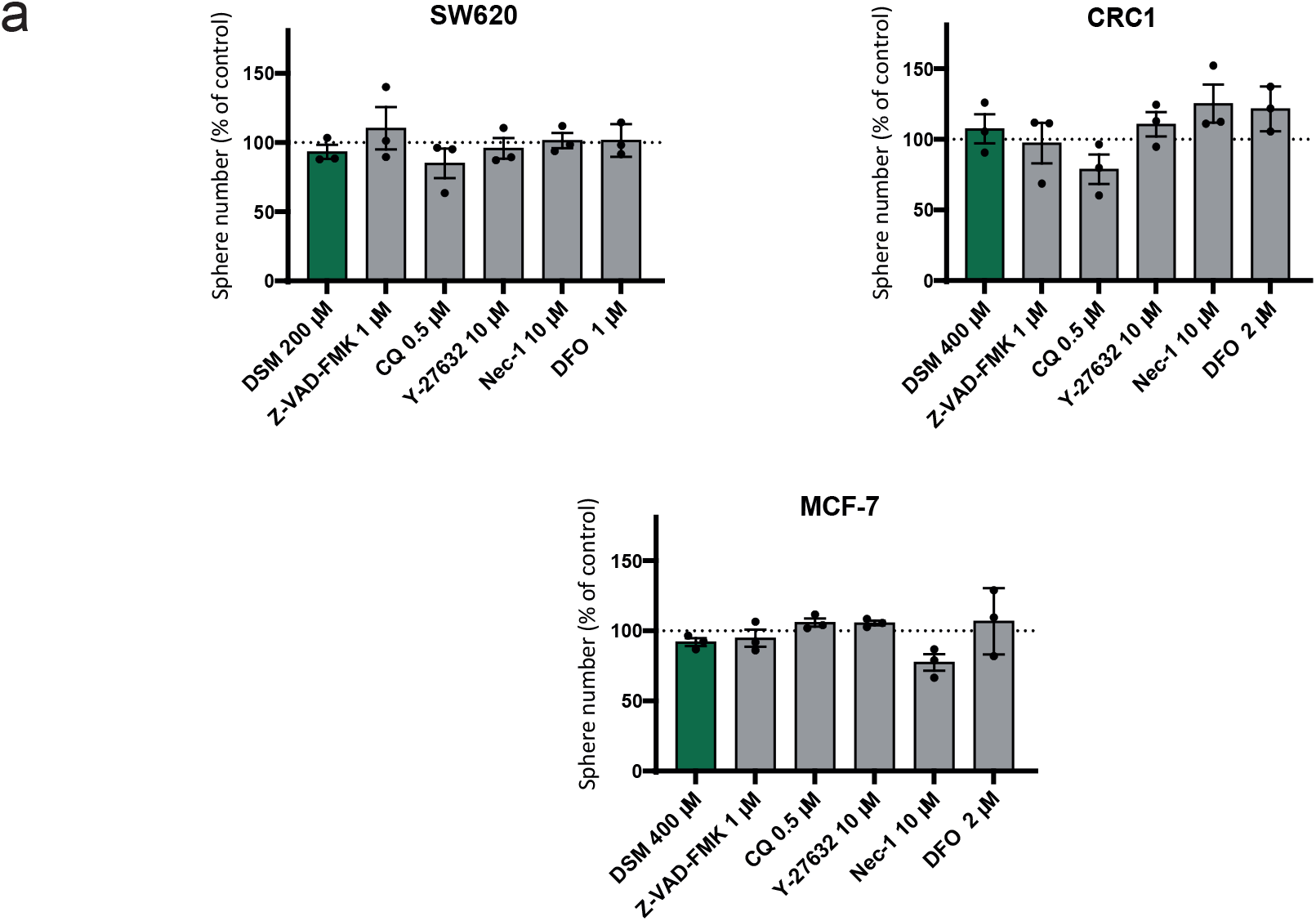
**a**. Cell death inhibitors do not affect sphere forming ability when used *in isolation.* Effects of Z-VAD-FMK (1 μM), CQ (0.5 μM), Y-27632 (10 μM), nec-1 (10 μM) and DFO (1 μM or 2 μM) added in SW620, CRC1 and MCF-7 cultured in suspension culture. Results are expressed in fold change compared to control. n=3 biological replicates. Mean ± SEM, ns sot significant. One way Anova followed by multiple comparisons.

**Figure S6.**
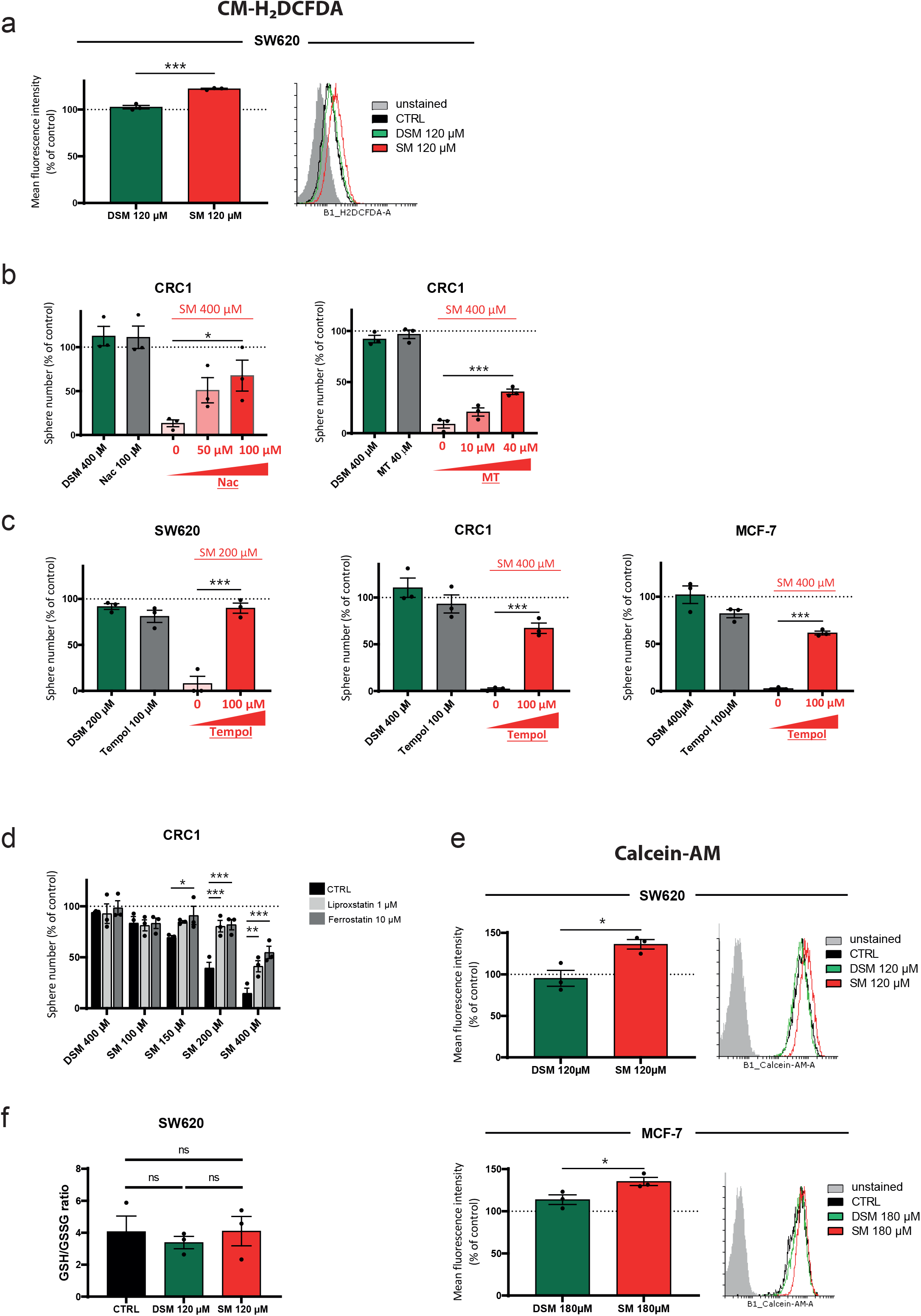
**a.** Total ROS level from SM-treated cells in suspension culture. Flow cytometry quantification of CM-H_2_DCFDA (50 nM SW620, 500 nM MCF-7) mean fluorescence intensity from suspension cultures of either DSM-or SM-treated cells (120 μM for SW620 or 180 μM for MCF-7). Results are expressed in fold change compared to control. n=3 biological replicates. Mean ± SEM, ns not significant, ***p-value < 0.001. Two-sided unpaired T-test. Histogram representative of triplicates. b. ROS inhibitors restore sphere formation of SM-treated CRC1 cells. SM-treated suspension culture of CRC1 (400 μM) were co-treated with Nac or MT at increasing concentrations. Results are expressed in fold change compared to control. n=3 biological replicates. Mean ± SEM, *p-value < 0.05, ***p-value < 0.001. One way Anova followed by multiple comparisons. c. The Superoxide dismutase mimetic Tempol inhibits SM effect on sphere formation. Cells treated with SM at 200 μM (SW620) or 400 μM (MCF-7 and CRC1) were co-treated with Tempol at 100 μM. Results are expressed in fold change compared to control. n=3 biological replicates. Mean ± SEM, ***p-value < 0.001. One way Anova followed by multiple comparisons. d. Lipid ROS inhibitors restore sphere formation in CRC1. CRC1 were pretreated in monolayer culture with Lip-1 (1 μM) or Fer-1 (10 μM). After 3 days, cells were passaged in suspension with increasing concentrations of SM. Results are expressed in fold change compared to control. n=3 biological replicates. Mean ± SEM, *p-value < 0.05, **p-value < 0.01, ***p-value < 0.001. Two way Anova followed by multiple comparisons. e. Cytoplasmic iron pool is increased following SM treatment in suspension culture. Flow cytometry quantification of Calcein-AM (10 nM) mean fluorescence intensity in cells cultured in suspension culture with DSM and SM at 120 μM (SW620) or 180 μM (MCF-7). Results are expressed in fold change compared to control. n=3 biological replicates. Mean ± SEM, *p-value < 0.05. Two-sided unpaired T-test. Histogram representative of triplicates. f. SM treatment does not impact GSH/GSSG ratio in suspension culture. GSH/GSSG ratio was calculated from SM- or DSM-treated (120 μM) SW620 cells grown in suspension for 6 days. Results are expressed in fold change compared to control. n=3 biological replicates. Mean ± SEM, ns not significant. One way Anova followed by multiple comparisons.

## Supplementary Tables

**Supplementary Table 1:**
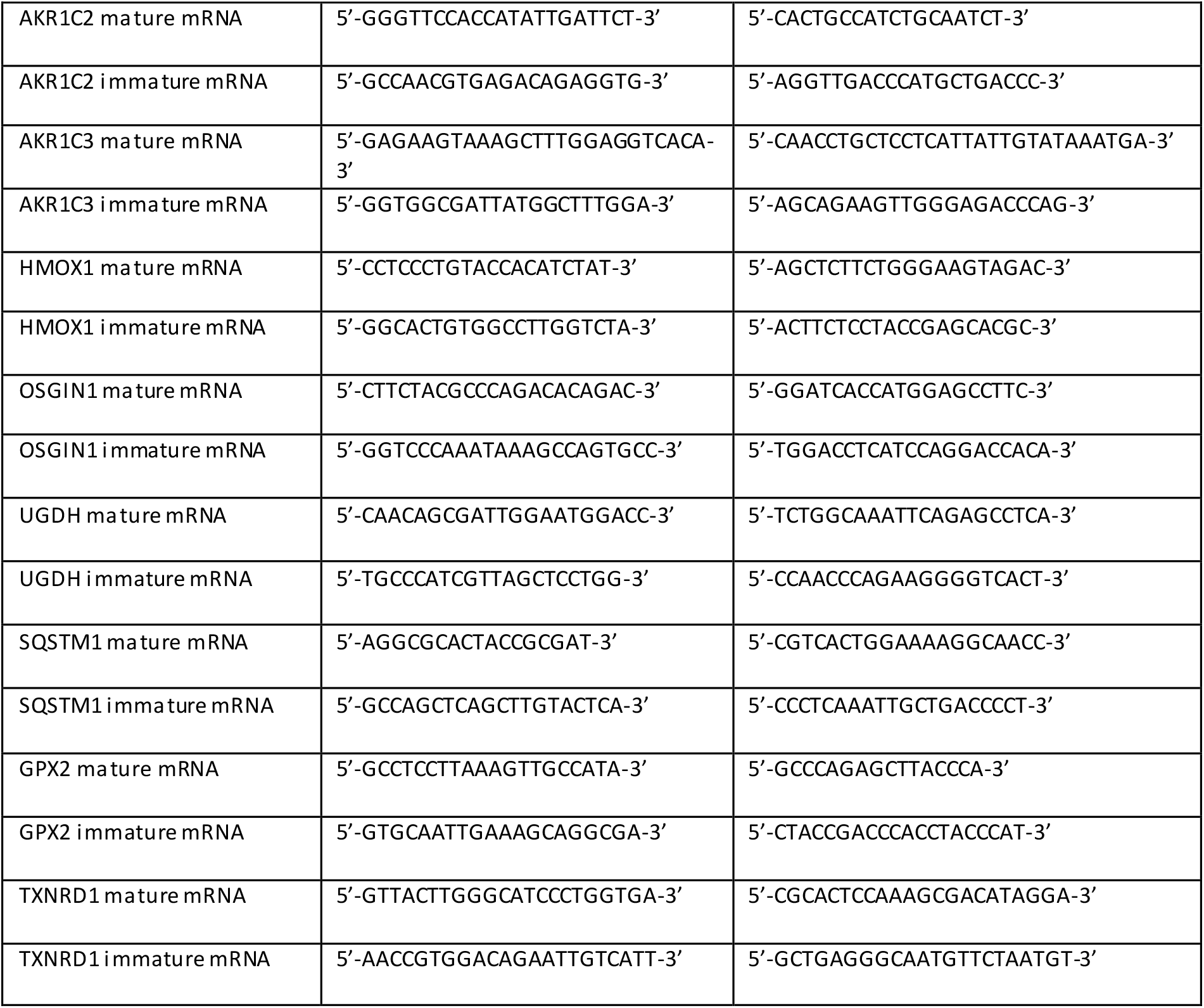
Primers used for the RNA-Seq validation.

**Supplementary Table 2:**
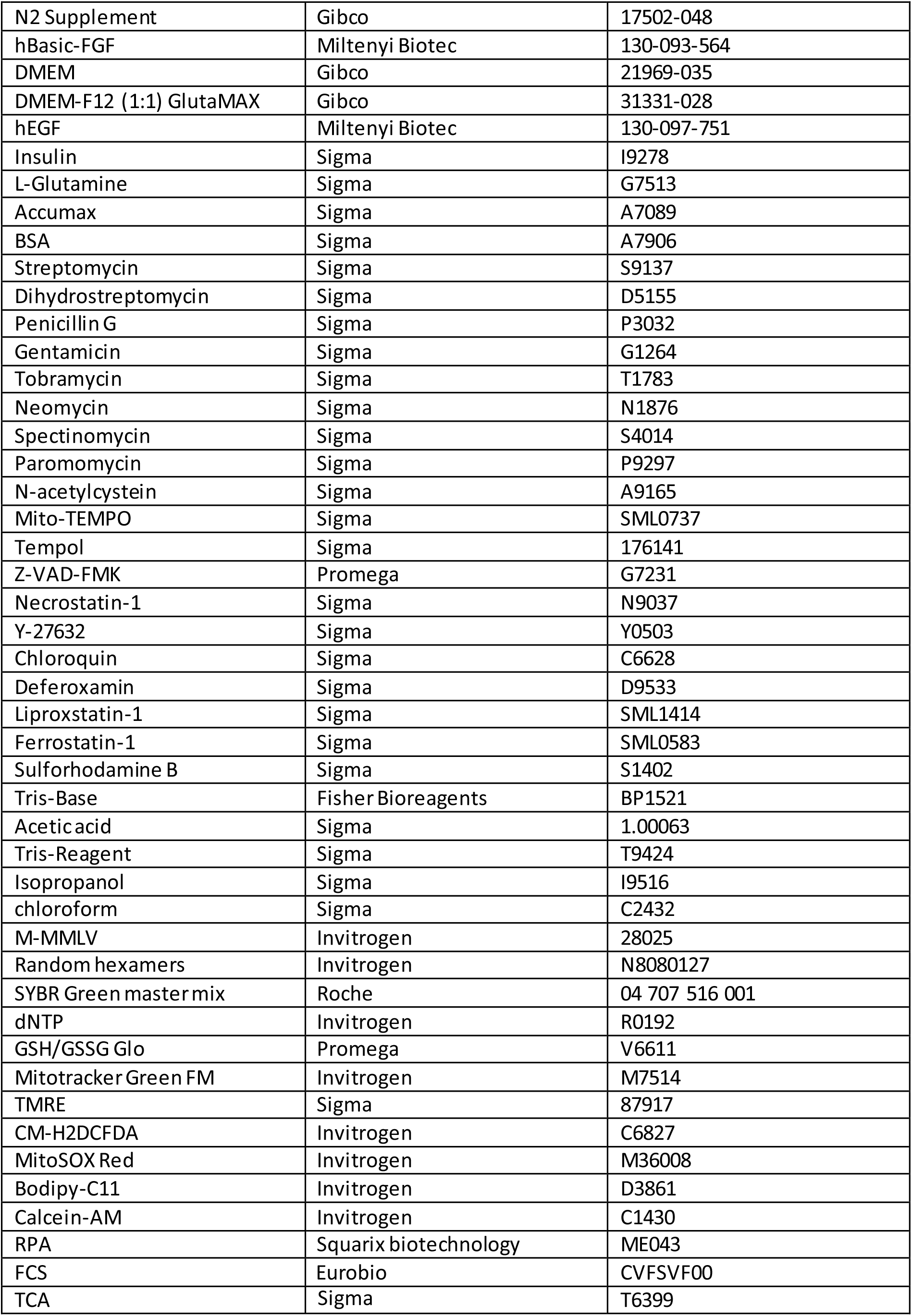
Reagents used in this work include.

## Chemistry: Supplementary material

Reagents and solvents were purchased from Merck and Carlo Erba Reagents and used without further purification. Streptomycin and dihydrostreptomycin were purchased from Merck as sulfate salts. HPLC was performed using a Water Alliance 2695 pump coupled with Waters 996 photodiode array detector and a Phenomenex Gemini® (5 μm, NX-C18, 4.6×250 mm Column for analytical HPLC and 10 × 150 mm for semipreparative HPLC). All HPLC analyses were run at room temperature using a gradient of CH_3_CN containing 0.1% TFA (eluent B) and water containing 0.1% TFA (eluent A) from 5% to 100% of B in 30 min at a flow rate of 1 mL/min for the analytical column and 3.5 mL/min for the semipreparative column. ^1^H and ^13^C NMR spectra were recorded on a Bruker AC 400 MHz spectrometer. Chemical shifts are reported in parts per million (ppm, δ) referenced to the residual ^1^H resonance of the solvent (D_2_O, δ 4.79). Splitting patterns are designated as follows: s (singlet), d (doublet), m (multiplet). Coupling constants (J values) are listed in hertz (Hz). Low resolution mass spectra (MS) were obtained with a Brucker Daltonics Esquire 3000+ electronspray spectrometer equipped with API ionization source. High resolution mass spectra (HRMS) were obtained with a LTQ Orbitrap hybrid mass spectrometer with an electronspray ionization probe (Thermoscientific, San Jose, CA) by direct infusion from a pump syringe to confirm correct molar mass and high purity of compounds.

## Synthesis Procedures

### Synthesis of streptomycin analogs: cf Figure S2

#### Synthesis of streptomycinic acid

Streptomycinic acid was synthesized starting from commercially available streptomycin sulfate (100 mg, 0.137 mmol) in one step upon treatment with Br_2_ (0.02 mL, 0.388 mmol, 2.8 eq.) in H_2_O (1.5 mL). The reaction mixture was stirred 5 days at room temperature and in the dark, then quenched using sodium thiosulfate. After filtration, the obtained filtrate was concentrated under reduced pressure and purified by semipreparative HPLC leading to desired streptomycinic acid in 73% yield (59.6 mg). Retention time 5.3 min;); ^1^H NMR (400 MHz, D_2_O) δ = 5.30-5.20 (m, 2H), 4.60-4.50 (m, 2H), 3.90-3.80 (m, 2H), 3.75-3.65 (m, 2H), 3.50-3.30 (m, 7H), 3.30-3.20 (m, 1H), 2.72 (s, 3H), 1.12 (d, J = 5.0 Hz, 3H); ^13^C NMR (125 MHz, D_2_O) d = 173.6, 158.1, 99.4, 93.5, 86.3, 81.4, 78.8, 74.7, 72.5, 71.5, 70.8, 69.3, 68.8, 60.5, 59.6, 58.9, 30.8, 12.2; HRMS (ESI): *m/z* 598.26807 (M+H)+ (C_21_H_41_O_13_N_7_ requires 598.26786).

### Synthesis of streptidine and streptobiosamine

Commercially available streptomycin was hydrolyzed to streptidine and streptobiosamine following a previously reported procedure (J. Biol. Chem. 1966, 241, 3142). Streptomycin (100 mg, 0.137 mmol) was dissolved in 1N sulfuric acid (1 mL) and stirred 48h at 37°C. The reaction mixture was then left 24h at 4°C to obtain a colorless precipitate of streptidine sulfate in 33% yield (20.7 mg). Retention time 7.7 min; ^1^H NMR (400 MHz, D_2_O) δ = 3.50-3.35 (m, 4H), 2.90-2.80 (m, 1H), 1.30-1.20 (m, 1H); MS (ESI): *m/z* 263.14624 (M+H)+ (C_8_H_20_O_4_N_6_ requires 263.14623). Acetone (1 mL) was then added to the filtrate and the solution left 24h at 4°C to precipitate the remaining streptidine. Lyophilization of the filtrate led to the obtention of streptobiosamine as a colorless solid in 27% yield (12.5 mg). Retention time 6.5 min; ^1^H NMR (400 MHz, D_2_O) δ = 5.46 (d, J = 2.0 Hz, 1H), 5.00-4.95 (m, 1H), 4.40-4.30 (m, 2H), 3.90-3.75 (m, 4H), 3.60-3.40 (m, 2H), 3.30-3.20 (m, 1H), 2.81 (s, 3H), 1.16 (d, J = 5.0 Hz, 3H); MS (ESI): *m/z* 370.17050 (M+MeOH)+ (C14H28NO10 370.17132).

**Figure S7.**
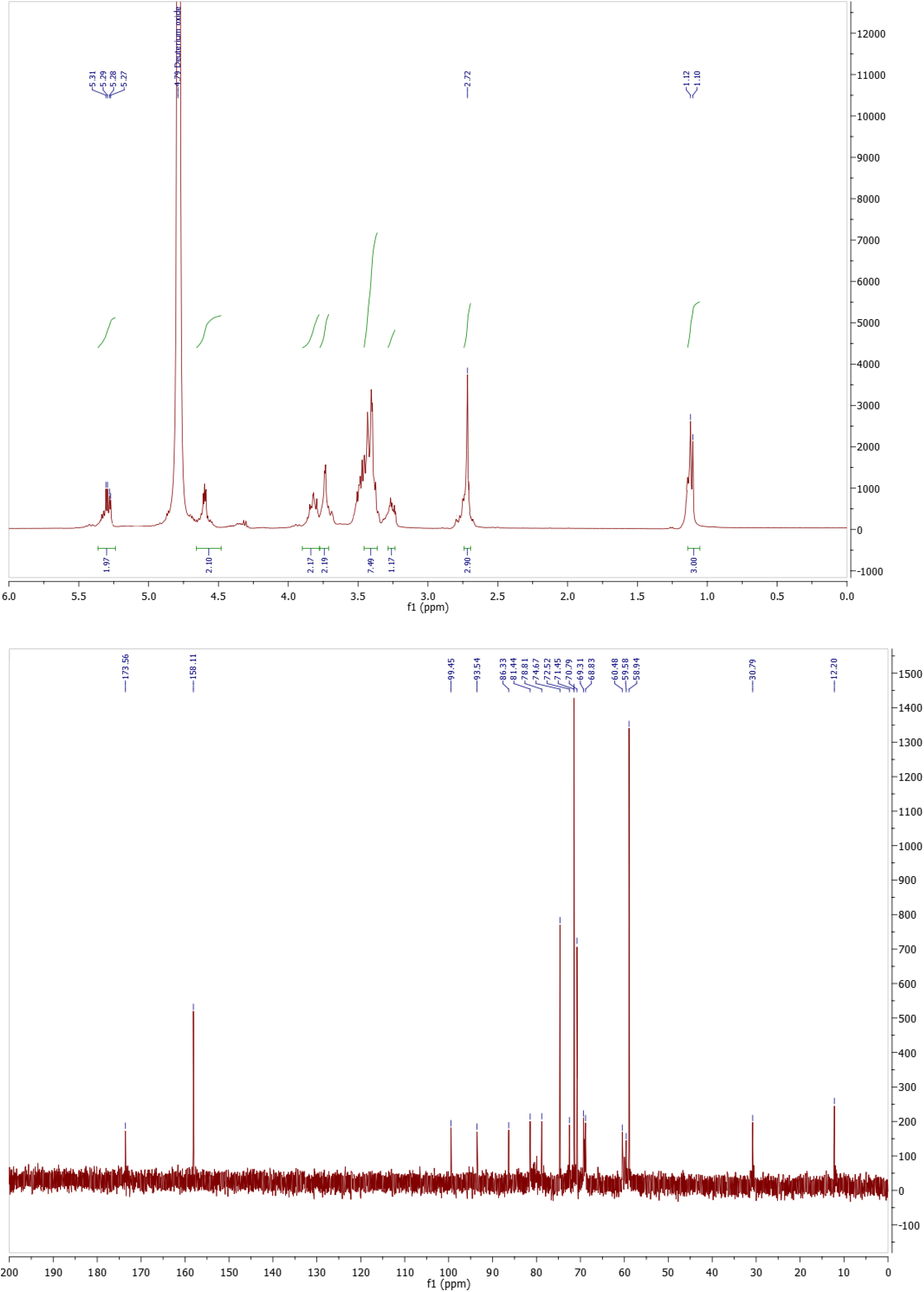
^1^H, ^13^CNMR and HRMS spectra for streptomycinicacid.

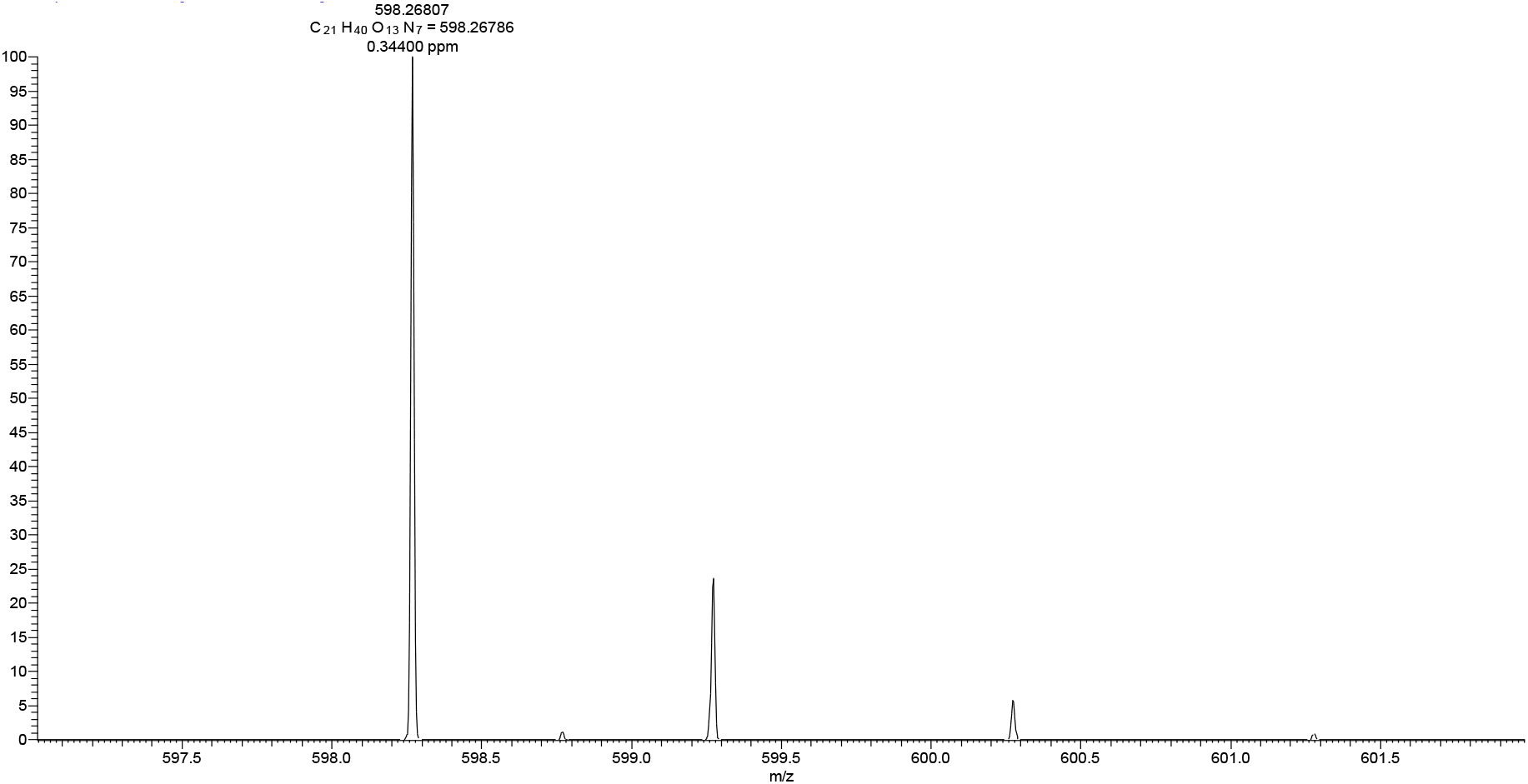

.1. J N, T Ö, K L, P B, Ml V. Methods for simultaneous and quantitative isolation of mitochondrial DNA, nuclear DNA and RNA from mammalian cells. BioTechniques [Internet]. déc 2020 [cité 15 juill 2021];69(6). Disponible sur: https://pubmed.ncbi.nlm.nih.gov/33103926/

